# Ventral tegmental area interneurons revisited: GABA and glutamate projection neurons make local synapses

**DOI:** 10.1101/2024.06.07.597996

**Authors:** Lucie Oriol, Melody Chao, Grace J. Kollman, Dina S. Dowlat, Sarthak M. Singhal, Thomas Steinkellner, Thomas S. Hnasko

## Abstract

The ventral tegmental area (VTA) contains projection neurons that release the neurotransmitters dopamine, GABA, and/or glutamate from distal synapses. VTA also contains GABA neurons that synapse locally on to dopamine neurons, synapses widely credited to a population of so-called VTA interneurons. Interneurons in cortex, striatum, and elsewhere have well-defined morphological features, physiological properties, and molecular markers, but such features have not been clearly described in VTA. Indeed, there is scant evidence that local and distal synapses originate from separate populations of VTA GABA neurons. In this study we tested whether several markers expressed in non-dopamine VTA neurons are selective markers of interneurons, defined as neurons that synapse locally but not distally. Challenging previous assumptions, we found that VTA neurons genetically defined by expression of parvalbumin, somatostatin, neurotensin, or mu-opioid receptor project to known VTA targets including nucleus accumbens, ventral pallidum, lateral habenula, and prefrontal cortex. Moreover, we provide evidence that VTA GABA and glutamate projection neurons make functional inhibitory or excitatory synapses locally within VTA. These findings suggest that local collaterals of VTA projection neurons could mediate functions prior attributed to VTA interneurons. This study underscores the need for a refined understanding of VTA connectivity to explain how heterogeneous VTA circuits mediate diverse functions related to reward, motivation, or addiction.

**Significance statement:** GABA neurons in VTA are key regulators of VTA dopamine neurons and considered central to the mechanisms by which opioids and other drugs of abuse can induce addiction. Conventionally, these VTA GABA neurons are considered interneurons, but GABA projection neurons are also abundant in VTA, and it is unclear if these represent separate populations. We found that several markers enriched in non-dopamine neurons of VTA, including Mu-opioid receptor, are also expressed in projection neurons, and thus are not selective interneuron markers. Moreover, we found that VTA GABA and glutamate projection neurons collateralize within VTA where they make local synapses. These data challenge the notion of a VTA interneuron that synapses only within VTA and suggest that inhibitory projection neurons can serve functions previously attributed to VTA interneurons.

## INTRODUCTION

The ventral tegmental area (VTA) is a central component of the brain’s reward circuitry, and a common attribute of addictive drugs is their ability to increase dopamine release from VTA projections (Luscher, 2016; Nestler, 2005). The VTA projects to and receives inputs from many brain structures involved in reward-related behavior, including nucleus accumbens (NAc), ventral pallidum (VP), lateral habenula (LHb), and prefrontal cortex (PFC) (Fields et al., 2007; Morales & Margolis, 2017). The VTA is often simplified as a region containing dopamine (DA) projection neurons and inhibitory GABA ‘interneurons’ that regulate DA neurons (Johnson & North, 1992; Luscher & Malenka, 2011; Nestler, 2005). However, VTA neurons are highly heterogeneous. The VTA contains distinct populations of DA neurons that can be segregated by gene expression, projection target, and function (Azcorra et al., 2023; Poulin et al., 2020; Roeper, 2013). GABA-releasing VTA neurons make local intra-VTA synapses (Bayer & Pickel, 1991; Omelchenko & Sesack, 2009), but also project widely outside the VTA, including dense projections to LHb, VP, and VP-adjacent areas of basal forebrain (Kaufling et al., 2010; Oades & Halliday, 1987; Taylor et al., 2014). Glutamate neurons are also prevalent in VTA and overlap with other populations such that ∼25% of VTA glutamate neurons co-express a DA marker and ∼25% express a GABA marker (Conrad et al., 2024; Ma et al., 2023; Phillips et al., 2022). VTA glutamate neurons release glutamate locally within VTA and from distal axons in medial NAc, PFC, VP, LHb, and elsewhere (Dobi et al., 2010; Gorelova et al., 2012; Hnasko et al., 2012; Root et al., 2014; Taylor et al., 2014; Yamaguchi et al., 2011).

It is now understood that DA signals can induce or correlate with distinct behavioral responses depending on their projection targets (Azcorra et al., 2023; Badrinarayan et al., 2012; de Jong et al., 2019; Faget et al., 2024). This is true also for VTA GABA and glutamate neurons. For example, activating VTA GABA neurons either locally within VTA or from distal processes can drive behavioral avoidance, disrupt reward seeking, or modify opioid reinforcement (Corre et al., 2018; Root et al., 2020; Shields et al., 2021; Soden et al., 2020; Tan et al., 2012; van Zessen et al., 2012; Zhou et al., 2022). On the other hand, stimulation of VTA GABA projections to LHb can be rewarding (Lammel et al., 2015; Stamatakis et al., 2013). Likewise, stimulation of VTA glutamate neurons can drive robust positive reinforcement or behavioral avoidance depending on the projection target and behavioral assay (Root et al., 2018; Root et al., 2014; Wang et al., 2015; Yoo et al., 2016). These responses can depend also on the co-release of distinct transmitters. For example, activation of VTA glutamate projections to NAc drives positive reinforcement through the release of glutamate and avoidance via DA co-release (Warlow et al., 2024; Zell et al., 2020). Thus, VTA neurons can mediate approach or avoidance behaviors through their specific connectivity and neurotransmitter content, and understanding the circuit mechanisms regulating activity in diverse VTA cell types is crucial to understanding the mechanisms by which mesolimbic circuits control motivated behaviors.

Local intra-VTA GABA modulation of VTA output, particularly DA output, may underlie key aspects of behavioral reinforcement. For example, inhibitory inputs to VTA from lateral hypothalamus, bed nucleus of stria terminalis, or VP can drive positive reinforcement and approach behaviors through inhibition of VTA GABA neurons and disinhibition of VTA DA neurons (Faget et al., 2024; Nieh et al., 2015; Soden et al., 2020; Soden et al., 2023). VTA GABA circuits also appear to be critical for the generation of DA reward prediction error signals (Eshel et al., 2015; Keiflin & Janak, 2015). Moreover, drugs of abuse can induce rapid or plastic changes in DA signaling through mechanisms that depend on intra-VTA GABA transmission (Corre et al., 2018; Gomez et al., 2019; Luscher & Malenka, 2011; Ostroumov & Dani, 2018; Ting & van der Kooy, 2012). Indeed, the observation that Mu-opioid receptor agonists directly inhibit non-DA VTA neurons and produce disinhibitory effects on VTA DA neurons (Johnson & North, 1992) helped establish the notion of VTA interneurons into current models of VTA architecture.

Yet there is scant evidence for the existence of VTA GABA interneurons, defined as neurons that make synapses locally within VTA but that do not make distal connections. Interneurons as so defined in cortex, striatum and other brain regions have characteristic morphological features, physiological properties, and molecular markers (Markram et al., 2004; Pelkey et al., 2017; Tepper et al., 2010). However, no molecular or physiological feature has been described that can clearly distinguish VTA GABA interneurons from GABA projection neurons. Identifying a marker that selectively labels VTA interneurons would enable investigations into distinct roles for VTA interneurons and projection neurons (Bouarab et al., 2019; Paul et al., 2019).

In this study we first sought to test whether several genes that are expressed in a subset of VTA neurons may be selective for interneurons in VTA. We chose markers that are expressed in non-DA neurons, selectively label interneurons in other brain areas, and/or have been widely presumed to label VTA interneurons. We found that these markers labeled neurons that were primarily non-DA neurons, but that made projections to distinct VTA projection targets, and thus did not selectively label VTA interneurons. We thus sought to test the hypothesis that VTA GABA (or glutamate) projection neurons make intra-VTA collaterals. Indeed, we provide both anatomical and physiological evidence that VTA GABA neurons projecting to NAc, VP, or PFC make local synapses within VTA. This work challenges the presumption of GABA interneurons in VTA by providing direct evidence for an alternative model by which GABA projection neurons can regulate the activity of neighboring VTA cells.

## RESULTS

### PV, SST, MOR, and NTS are not selective interneuron markers in VTA

We selected four genes with well-validated Cre lines to test as putative genetic markers that might selectively label VTA interneurons: Parvalbumin (PV), Somatostatin (SST), Mu-opioid receptor (MOR), or Neurotensin (NTS). We injected adeno-associated virus (AAV) into the VTA for Cre-dependent expression of Channelrhodopsin-2 (ChR2) fused to mCherry that labels distal axons (**Figure 1A**). To test whether the labeled VTA neurons project distally we assessed expression in known VTA projection sites including nucleus accumbens (NAc), ventral pallidum (VP), prefrontal cortex (PFC), and lateral habenula (LHb).

**Figure 1:**
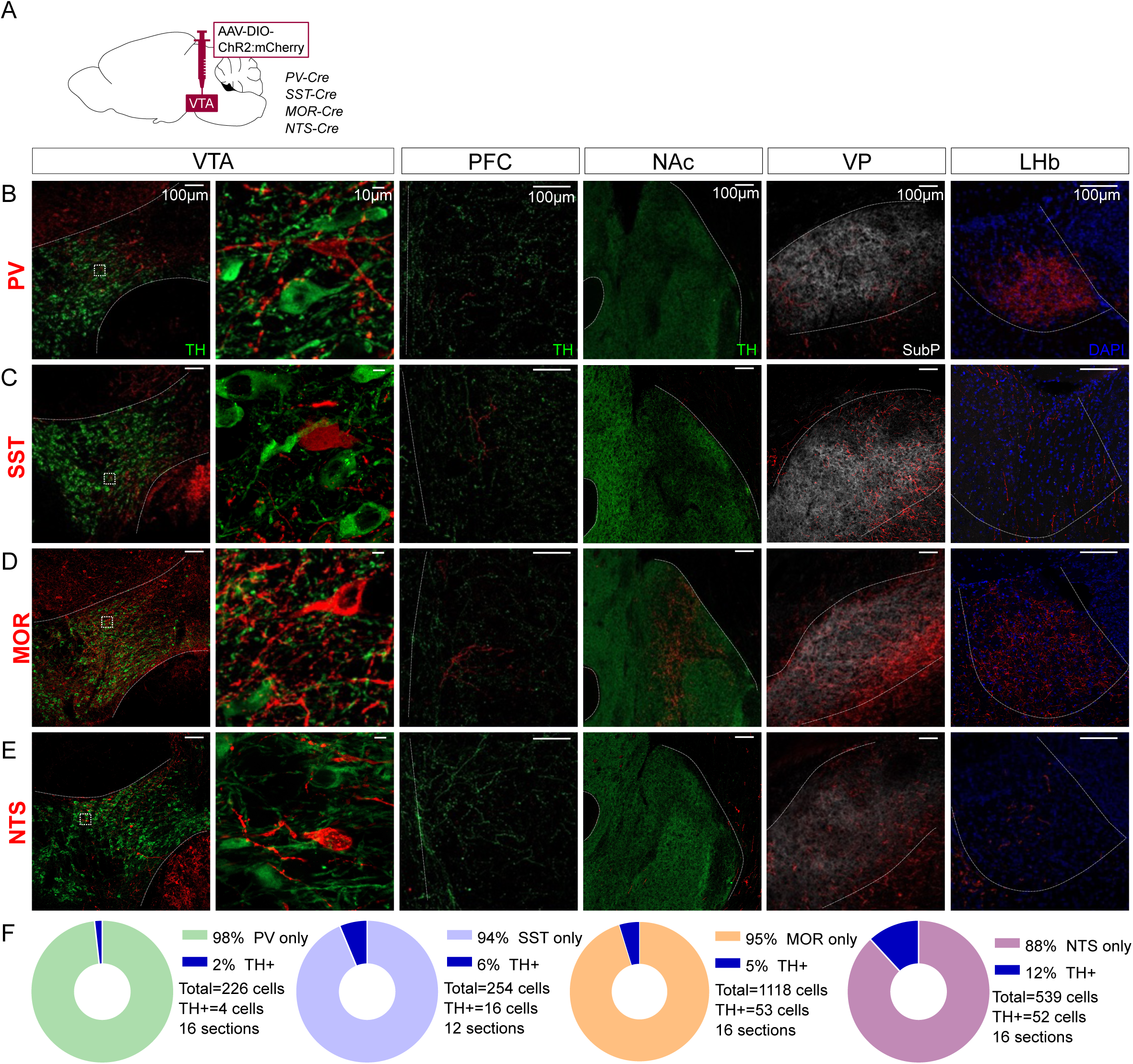
Distal projections of putative VTA interneuron markers. (**A**) Cre-dependent expression of ChR2:mCherry in VTA cell bodies but also distal axonal process in (**B**) PV-Cre, (**C**) SST-Cre, (**D**) MOR-Cre, and (**E**) NTS-Cre mice. First column is an overview of the expression in VTA (bregma -3.3), followed by a high magnification inset of the boxed region in the second column. The third column shows expression patterns in PFC (bregma +1.7), the fourth in NAc (bregma +1.3), the fifth in VP (bregma +0.5) and the sixth in LHb (bregma -1.8). Scale bars are 100µm, except 10µm in the second column. ChR2:mCherry is shown in red; with TH in green, Substance P in white, or DAPI in blue. (**F**) Donut charts show the fraction of mCherry+ VTA cells counted that label for TH.

Parvalbumin (PV) is a marker of interneurons in cortex and striatum (Kawaguchi, 1993; Kawaguchi & Kondo, 2002; Tepper et al., 2018), but is also expressed in VTA GABA neurons (Olson & Nestler, 2007). Injections into VTA of PV-Cre mice labeled neurons located in medial VTA. We also detected a dense concentration of axonal fibers in LHb, with scant labeling in other known VTA projection targets (**Figure 1B**). These data suggest that PV labels VTA projection neurons and PV is not a selective marker of VTA interneurons.

Like PV, Somatostatin (SST) is an interneuron marker in cortex (Kawaguchi & Kondo, 2002). SST is expressed in VTA GABA neurons that can inhibit neighboring VTA DA neurons (Nagaeva et al., 2020). Injections into SST-Cre mice labeled cell bodies in VTA (**Figure 1C**). We again identified axons in distal targets, here with notably dense labeling in VP.

MOR is expressed in VTA GABA neurons, inhibiting GABA release from synapses on to VTA DA neurons, thereby increasing DA neuron firing, and is often described as a marker of VTA interneurons (Gysling & Wang, 1983; Johnson & North, 1992; Luscher & Malenka, 2011; Nestler, 2005; Phillips et al., 2022). Injections into MOR-Cre mice led to labeled neurons throughout VTA, but also labeled axons in PFC, NAc, LHb, and especially VP (**Figure 1D**).

NTS is expressed in a subpopulation of VTA GABA neurons (Phillips et al., 2022) and NTS can stimulate mesolimbic DA cells through activation of NTS receptor 1 (Caceda et al., 2006; Kalivas, 1983). Injections into NTS-Cre mice labeled neurons in VTA, as well as axons in VP, with weaker labeling in other VTA projection sites (**Figure 1E**).

We also stained VTA sections for Tyrosine hydroxylase (TH) to estimate the proportion of ChR2:mCherry neurons colocalizing with DA neurons. In all cases only a minority of mCherry-labeled neurons expressed TH, ranging from 2% for PV to 12% for NTS (**Figure 1F**). In total, our data suggest that these four markers label primarily non-DA neurons in VTA, but that none are selective for interneurons, and instead are inclusive of VTA projection neurons.

### Anatomical evidence that VTA projection neurons make local synapses

Each of the markers tested are also expressed in neurons proximal to VTA and our injections led to variable spread to neighboring regions, including interpeduncular nucleus (IPN) and red nucleus. While these regions are not known to project to PFC, NAc, VP, or LHb, we nonetheless aimed to validate the above findings with a secondary approach involving a combination of retrograde labeling and intersectional genetics to target VTA projection neurons. We injected AAV-fDIO-mGFP-Synaptophysin:mRuby into VTA of each Cre line, plus retroAAV-DIO-Flp into a projection target receiving dense innervation. This approach allowed for Cre-plus Flp-dependent expression of both membrane-localized GFP and the synaptic vesicle marker Syn:Ruby (Beier et al., 2015). The intersectionality of this approach allows for precise targeting of VTA projection neurons, and Syn:Ruby highlights putative release sites, either local to or distal from VTA.

Using this approach to label PV-Cre projectors to LHb (**Figure 2A**), or SST-Cre projectors to VP (**Figure 2G**), revealed GFP-positive soma well-restricted within VTA borders delineated by TH immunolabel (**Figure 2B,C,H,I**). We also observed GFP-positive axons and Syn:Ruby-positive puncta in LHb of PV-Cre, or VP of SST-Cre mice (**Figure 2E,F,K,L**). Using high magnification we observed Syn:Ruby puncta proximal to TH-positive cells in VTA (**Figure 2D,J**), suggesting that these VTA projection neurons collateralize within VTA and synapse on to DA neurons.

**Figure 2:**
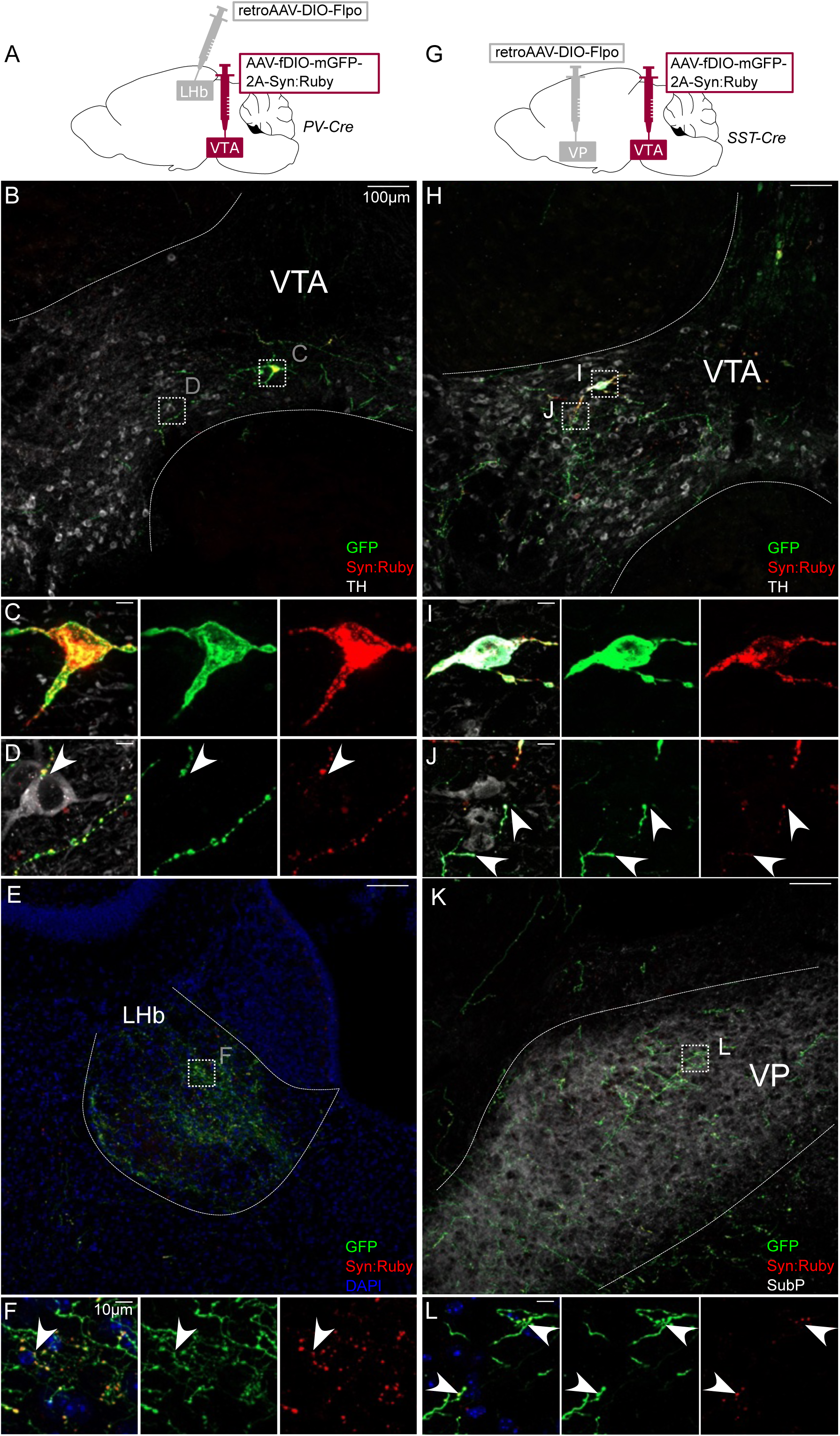
Intersectional approach to label projections of PV- and SST-expressing VTA neurons. (**A**) Dual AAV approach for Cre-dependent expression of Flp injected in LHb plus Flp-dependent expression of GFP and Syn:Ruby in VTA of PV-Cre mice. (**B**) LHb-projecting PV-Cre neurons in VTA with (**C,D**) high magnification insets showing putative release sites proximal to TH+ DA neurons. (**E**) VTA axons in LHb with (**F**) high magnification insets. (**G**) Dual AAV approach for Cre-dependent expression of Flp injected in VP plus Flp-dependent expression of GFP and Syn:Ruby in VTA of SST-Cre mice. (**H**) VP-projecting SST-Cre neurons in VTA with (**I,J**) high magnification insets showing putative release sites proximal to TH+ DA neurons. (**K**) VTA axons in VP with (**L**) high magnification insets. Scale bars 100µm, or 10µm for high magnification insets.

The same approach was used to label VTA projectors to VP in MOR-Cre or NTS-Cre mice (**Figure 3A**). VP-projecting GFP-positive cell bodies in MOR-Cre mice were contained within and throughout VTA (**Figure 3B**). We also observed axonal fibers densely filling VP, delineated by Substance P immunolabel (**Figure 3F-H and Supplemental Figure 1**). Using high magnification, we observed Syn:Ruby puncta proximal to TH-positive VTA neurons, again suggestive of synapses made within VTA by labeled projection neurons (**Figure 3C-E**). Using the same approach to target VP-projecting neurons in NTS-Cre mice we found similar results, with GFP-positive soma in VTA and Syn:Ruby-positive puncta in both VTA (**Figure 3I-J**) and VP (**Figure 3K-L**), though signals were notably less dense. These results corroborate the findings in **Figure 1** and suggest that multiple markers that had been suggested to label putative interneurons instead label VTA projection neurons that may make local synapses through axon collaterals.

**Figure 3:**
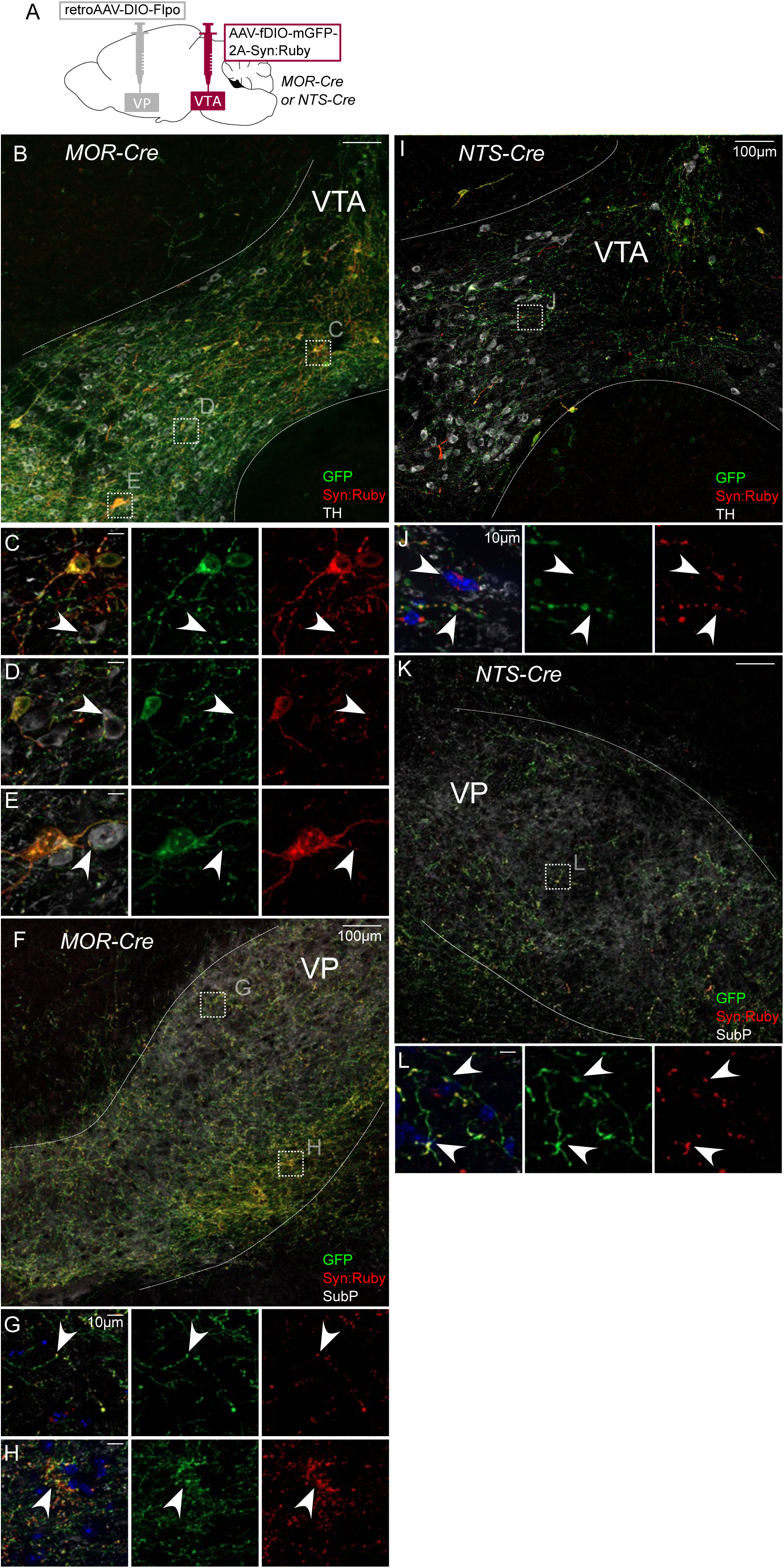
Intersectional approach to label projections of MOR- and NTS-expressing VTA neurons. (**A**) Dual AAV approach for Cre-dependent expression of Flp injected in VP plus Flp-dependent expression of GFP and Syn:Ruby in VTA of MOR-Cre and NTS-Cre mice. (**B**) VP-projecting MOR-Cre neurons in VTA with (**C,D,E**) high magnification insets showing putative release sites proximal to TH+ DA neurons. (**F**) VTA axons in VP with (**G,H**) high magnification insets. (**I**) VP-projecting NTS-Cre neurons in VTA with (**J**) high magnification insets showing putative release sites. (**K**) VTA axons in VP with (**L**) high magnification insets. Scale bars 100µm, or 10µm for high magnification insets.

There is evidence indicating that PV, SST, MOR and NTS neurons in VTA express GABA markers or release GABA (Nagaeva et al., 2020; Olson & Nestler, 2007; Phillips et al., 2022). However, some neurons positive for those markers may express VGLUT2 and release glutamate (Miranda-Barrientos et al., 2021). We therefore used VGAT-Cre and VGLUT2-Cre mice to selectively express GFP and Syn:Ruby, here targeting NAc-projecting VTA neurons. In VGAT-Cre mice (**Figure 4A**) we identified GFP-positive cell bodies that were restricted to VTA (**Figure 4B**) and GFP-positive fibers in NAc (**Figure 4E-F**). At higher magnification, we observed a pattern of GFP fibers and Syn:Ruby puncta surrounding TH-positive cell bodies (**Figure 4C-D**), suggesting that NAc-projectors make collaterals on to VTA DA neurons. We observed similar results when using VGLUT2-Cre mice, suggesting that NAc-projecting VTA glutamate neurons can also make local collaterals within VTA (**Figure 4G-L**). As expected VTA glutamate cell bodies were concentrated in medial VTA, where they are most dense (Conrad et al., 2024; Kawano et al., 2006; Yamaguchi et al., 2011).

**Figure 4:**
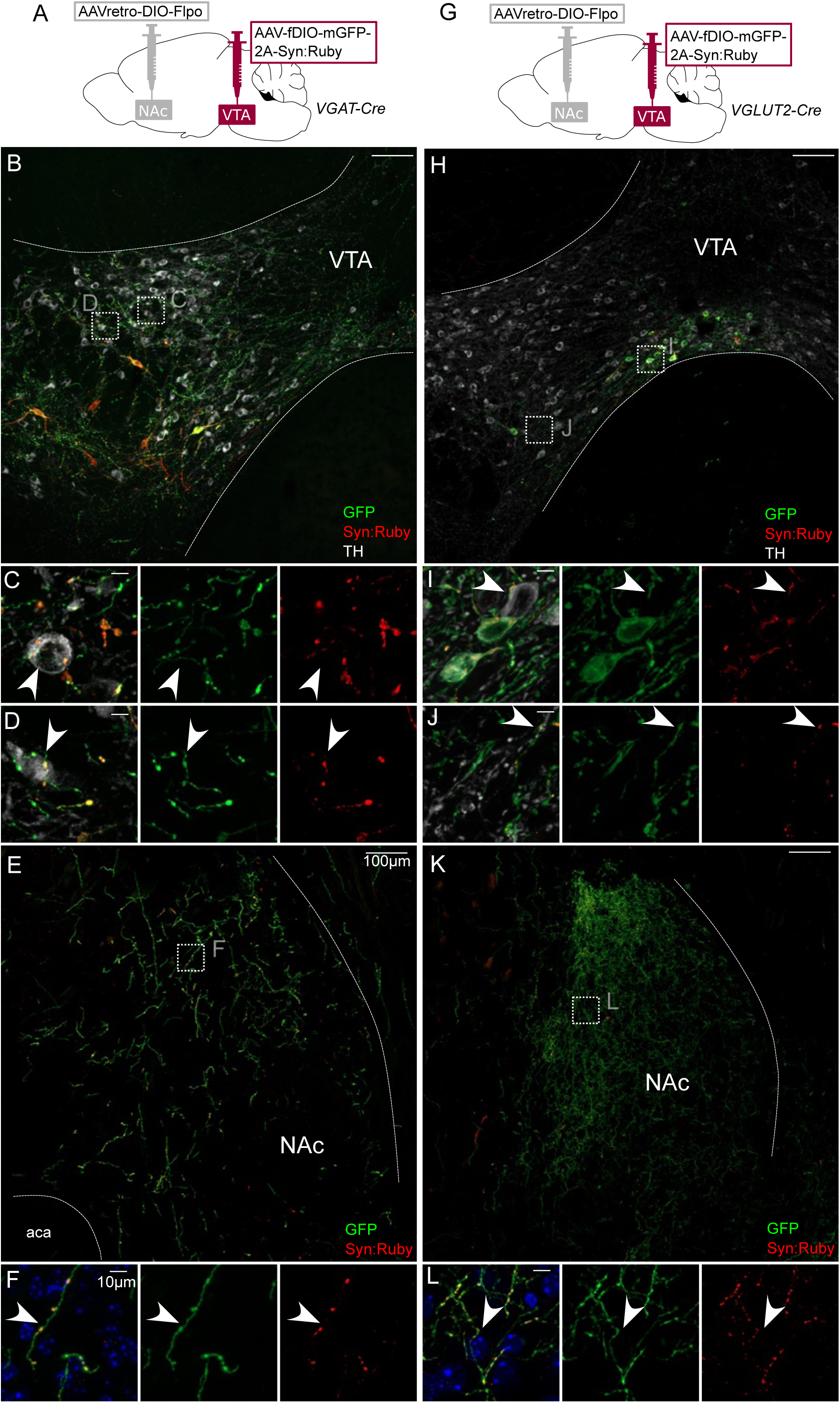
Intersectional labeling of VTA GABA and glutamate projection neurons suggests intra-VTA collaterals. (**A**) Dual AAV approach for Cre-dependent expression of Flp injected in NAc plus Flp-dependent expression of GFP and Syn:Ruby in VTA of VGAT-Cre mice. (**B**) NAc-projecting VGAT-Cre neurons in VTA with (**C,D**) high magnification insets showing putative release sites proximal to TH+ DA neurons. (**E**) VTA axons in NAc of VGAT-Cre mice, with (**F**) high magnification insets. (**G**) Dual AAV approach for Cre-dependent expression of Flp injected in NAc plus Flp-dependent expression of GFP and Syn:Ruby in VTA of VGLUT2-Cre mice. (**H**) NAc-projecting VGLUT2-Cre neurons in VTA with (**I,J**) high magnification insets showing putative release sites proximal to TH+ DA neurons. (**K**) VTA axons in NAc of VGLUT2-Cre mice with (**L**) high magnification insets. Scale bars 100µm, or 10µm for high magnification insets.

### Physiological evidence that VTA projection neurons make local synapses

Our anatomical results suggest that multiple types of VTA projection neurons collateralize locally within VTA. Next, to functionally assess whether VTA projection neurons make local synapses in VTA, we used a combination of optogenetics and electrophysiology. We selectively expressed ChR2 in NAc-projecting VTA neurons by injecting retroAAV-Cre into NAc and AAV-DIO-ChR2:mCherry into VTA of wild-type mice (**Figure 5A**). We then made acute brain slices to record from VTA neurons negative for ChR2:mcherry to test if they received synaptic inputs from NAc-projecting VTA neurons (**Figure 5B**). Using wild-type mice allowed us to express opsin in both GABA and glutamate projection neurons, and assess for optogenetic-evoked postsynaptic currents (oPSC) that were either inhibitory (oIPSC) or excitatory (oEPSC) from the same cell.

**Figure 5:**
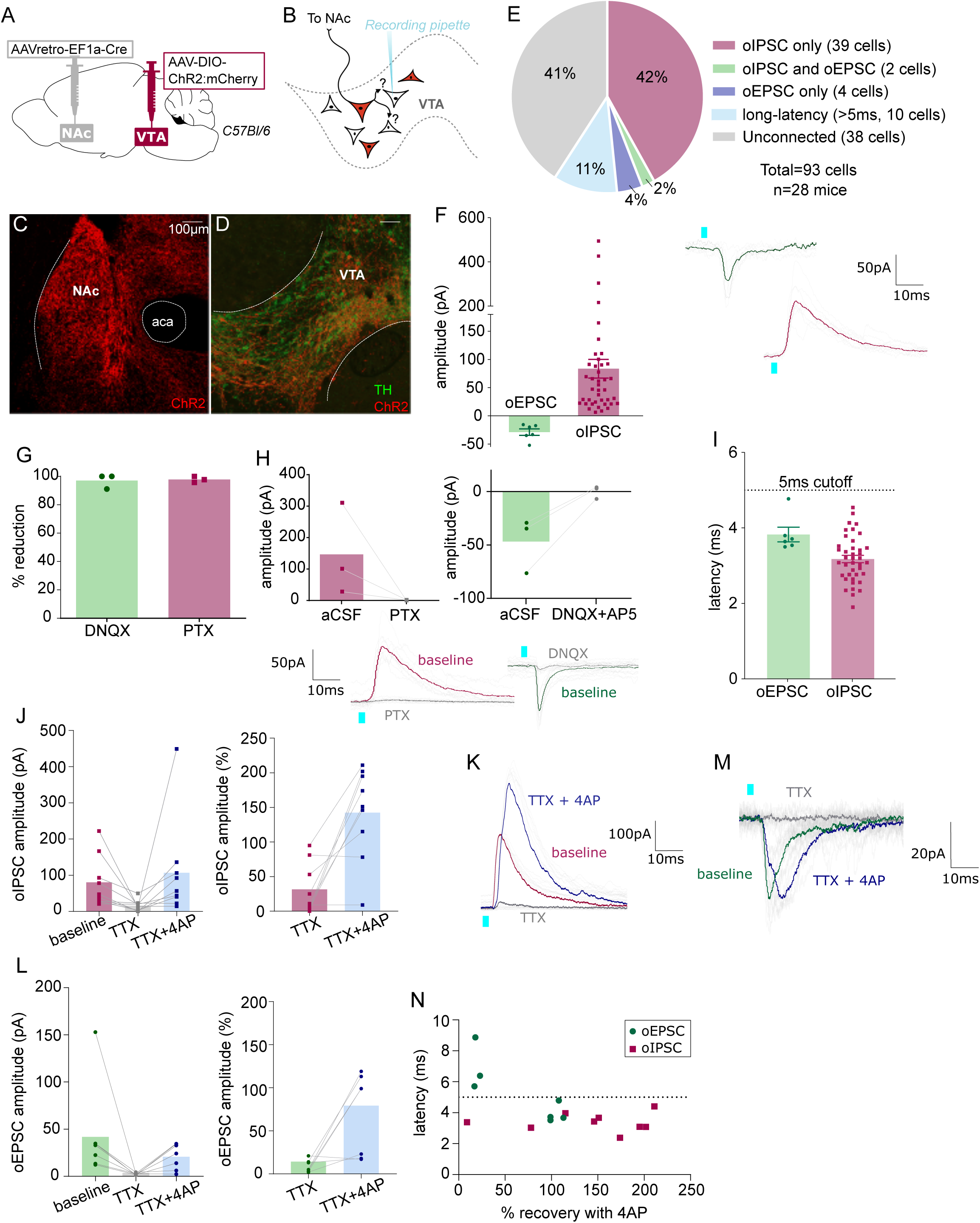
NAc-projecting VTA GABA and glutamate neurons make intra-VTA synapses. (**A**) Dual AAV approach to express ChR2:mCherry in NAc-projecting VTA neurons in wild-type mice. (**B**) Patch-clamp recordings from ChR2:mCherry-negative neurons of VTA to test for collateralizing synapses made by NAc-projectors. (**C**) Coronal images showing ChR2:mCherry expression in NAc and (**D**) VTA; scale bars 100 µm. (**E**) ChR2:mCherry-negative VTA neuron responses to optogenetic stimulation of NAc-projectors. (**F**) Peak amplitude of connected cells that displayed an oEPSC and/or oIPSC (excluding long-latency), with example traces. (**G**) Percent reduction in oEPSC or oIPSC by DNQX or PTX, respectively. (**H**) Peak amplitude of oIPSCs before and after bath application of PTX, or of oEPSCs before and after bath application of DNQX, with example traces. (**I**) Latency to oPSC onset (excluding long-latency). (**J**) Peak oIPSC amplitude before and after bath application of TTX and recovery with 4AP (Friedman’s test Chi-square=10.9, p=0.0029) and (**K**) example traces. (**L**) Peak oEPSC amplitude before and after bath application of TTX and recovery with 4AP (Friedman’s test Chi-square=11.6, p=0.0013) and (**M**) example traces (**N**) Scatter plot showing relationship between initial (pre-treatment) latency to oPSC onset and 4AP recovery. Green dots represent oEPSCs and red squares oIPSCs.

As expected, the medial shell of NAc showed dense mCherry-positive fibers (**Figure 5C**), and mCherry-positive cell bodies were restricted to VTA (**Figure 5D**). We patched ChR2:mCherry-negative VTA neurons (**Supplemental Figure 2**), flashed 2-ms blue light pulses, and observed oPSCs in 59% of neurons; 44% displayed short-latency oIPSCs (mean 84 ± 17 pA), 6% had short-latency oEPSCs (mean -28 ± 6 pA), 41% had no response (responses less than 5 pA were considered unconnected), and 11% had oPSCs with long-latency to onset (>5 ms) (**Figure 5E-F**). Note that unconnected and long-latency cells are not included in **Figures 5F, I**. The GABAA receptor antagonist picrotoxin (PTX) blocked oIPSCs while oEPSCs were blocked by the AMPA receptor antagonist DNQX (**Figure 5G,H**), confirming that these responses are mediated by evoked GABA or glutamate release, respectively.

Most responses displayed onset latencies more than 2 ms and less than 5 ms, consistent with monosynaptic connectivity (3.2 ± 0.1 ms and 3.8 ± 0.2 ms for oIPSCs and oEPSCs, respectively) (**Figure 5I**). To confirm connections are monosynaptic we performed additional pharmacology. We found that the amplitude of oPSCs was diminished following the application of the voltage-gated sodium channel blocker tetrodotoxin (TTX, voltage-gated sodium channel are necessary for the propagation of action potentials), and that oPSCs recovered with bath application of the inhibitor of voltage-sensitive potassium channels 4-aminopyridine (4AP). When this strategy was applied to oIPSCs (**Figure 5J**), 8 out of 9 TTX-diminished currents were restored by the application of 4AP (**Figure 5J,K**). Similarly, 4 of 7 oEPSCs were recovered by 4AP (**Figure 5L,M**). We plotted the latency of oPSC onset against the percent oPSC recovery mediated by 4AP and found that 3 of 4 neurons that failed to recover had a latency >5ms, whereas only 1 of 13 neurons that had a latency <5 ms failed to recover (**Figure 5N**). Therefore, we used 5 ms as a ‘short latency’ cutoff to consider an oPSC as monosynaptic. In total we recorded ten cells with oPSC latency >5 ms (identified as long latency on **Figure 5E**). Out of these 10 long-latency oPSCs, 8 were oEPSCs and 2 oIPSCs. This proportion (8:2) of neurons with oEPSCs versus oIPSCs was strikingly greater than that for short-latency responses (6:41), suggesting that in some cells/slices optogenetic stimulation of projection neurons recruited a more extensive intra-VTA excitatory network.

We used a similar approach to assess whether VTA neurons projecting to VP or PFC also made local collaterals in VTA. We used the same combination of viruses but here injected retroAAV-Cre into VP of wild-type mice (**Figure 6A**), again recording from mCherry-negative VTA neurons (**Figure 6B**). As expected, we observed dense mCherry-positive fibers in VP and mCherry-positive cell bodies restricted to VTA (**Figure 6C-D and Supplemental Figure 3**). We found that 52% of mCherry-negative neurons were connected (13 of 25), 32% displayed short-latency oIPSCs, 4% had short-latency oEPSCs, and 20% were connected but with long latency (>5 ms) (**Figure 6E-G**). We also patched from postsynaptic neurons in VP and found 89% displayed oPSCs, all with short latency, and as in VTA most currents were inhibitory (**Figure 6H-K**).

**Figure 6:**
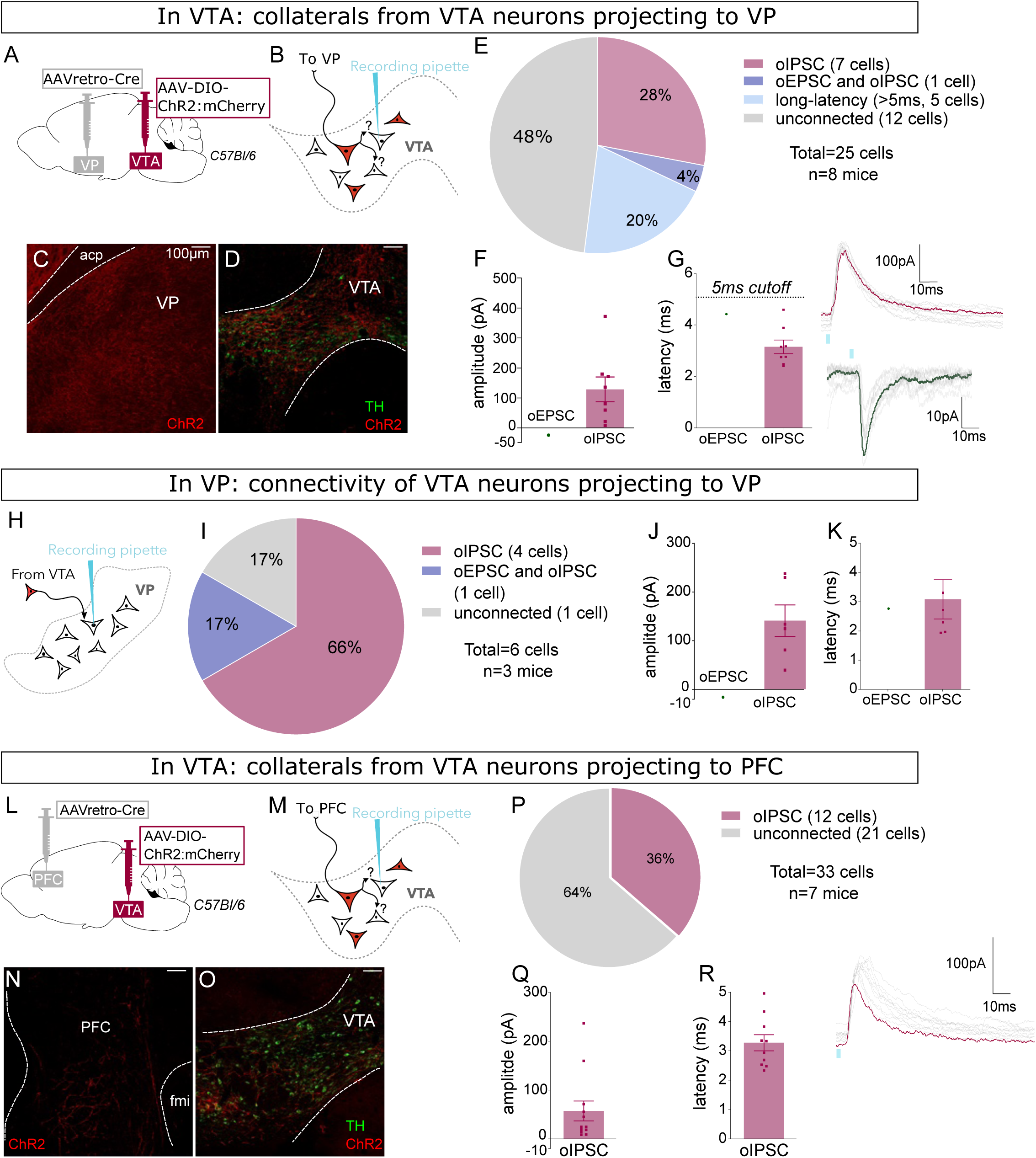
VP-projecting and PFC-projecting VTA GABA and glutamate neurons make intra-VTA synapses. (**A**) Dual AAV approach to express ChR2:mCherry in VP-projecting VTA neurons in wild-type mice. (**B**) Patch-clamp recordings from ChR2:mCherry-negative neurons of VTA to test for collateralizing synapses made by VP-projectors. (**C**) Coronal images showing ChR2:mCherry expression in VP and (**D**) VTA; scale bars 100 µm. (**E**) ChR2:mCherry-negative VTA neuron responses to optogenetic stimulation of VP-projectors. (**F**) Peak amplitude and (**G**) onset latency of connected cells that displayed an oEPSC and/or oIPSC (excluding long-latency), with example traces. (**H**) Recordings of oPSCs from neurons in VP and (**I**) VP responses to optogenetic stimulation of VP-projecting VTA neurons from approach described in panel A. (**J**) Peak amplitude and (**K**) onset latency of connected VP neurons that displayed an oEPSC and/or oIPSC, with example traces. (**L**) Dual AAV approach to express ChR2:mCherry in PFC-projecting VTA neurons in wild-type mice. (**M**) Patch-clamp recordings from ChR2:mCherry-negative neurons of VTA to test for collateralizing synapses made by PFC-projectors. (**N**) Coronal images showing ChR2:mCherry expression in PFC and (**O**) VTA; scale bars 100 µm. (**P**) ChR2:mCherry-negative VTA neuron responses to optogenetic stimulation of PFC-projectors. (**Q**) Peak amplitude and (**R**) onset latency of connected cells that displayed an oEPSC and/or oIPSC, with example trace.

Finally, we repeated the same approach but for PFC-projecting VTA neurons (**Figure 6L-M**). We observed mCherry-positive fibers in PFC (**Figure 6N**) arising from sparse cell bodies found within the bounds of VTA (**Figure 6O and Supplemental Figure 3**). In VTA, we found that 36% of ChR2:mCherry-negative neurons were connected, all of which displayed short-latency oIPSCs (**Figure 6P-R**). Altogether our data indicate that VTA GABAergic projection neurons, and to a lesser extent glutamatergic projection neurons, make functional synapses within VTA.

The use of WT mice in these experiments allowed us to assay for the presence of inhibitory currents mediated by GABA-releasing neurons and for excitatory currents mediated by glutamate-releasing neurons, from the same postsynaptic cells. However, we also performed similar experiments using VGAT-Cre and MOR-Cre mice. In these experiments we used an intersectional approach similar to **Figures 2-4**. We targeted NAc-projecting VGAT-Cre neurons by injecting retroAAV for Cre-dependent expression of Flp into NAc, plus AAV for Flp-dependent expression of ChR2:YFP into VTA (**Figure 7A-D**). We found that the majority of YFP-negative VTA neurons that we recorded displayed short-latency oIPSCs (**Figure 7E-G**) (note that we did not assay for oEPSCs in this experiment).

**Figure 7.**
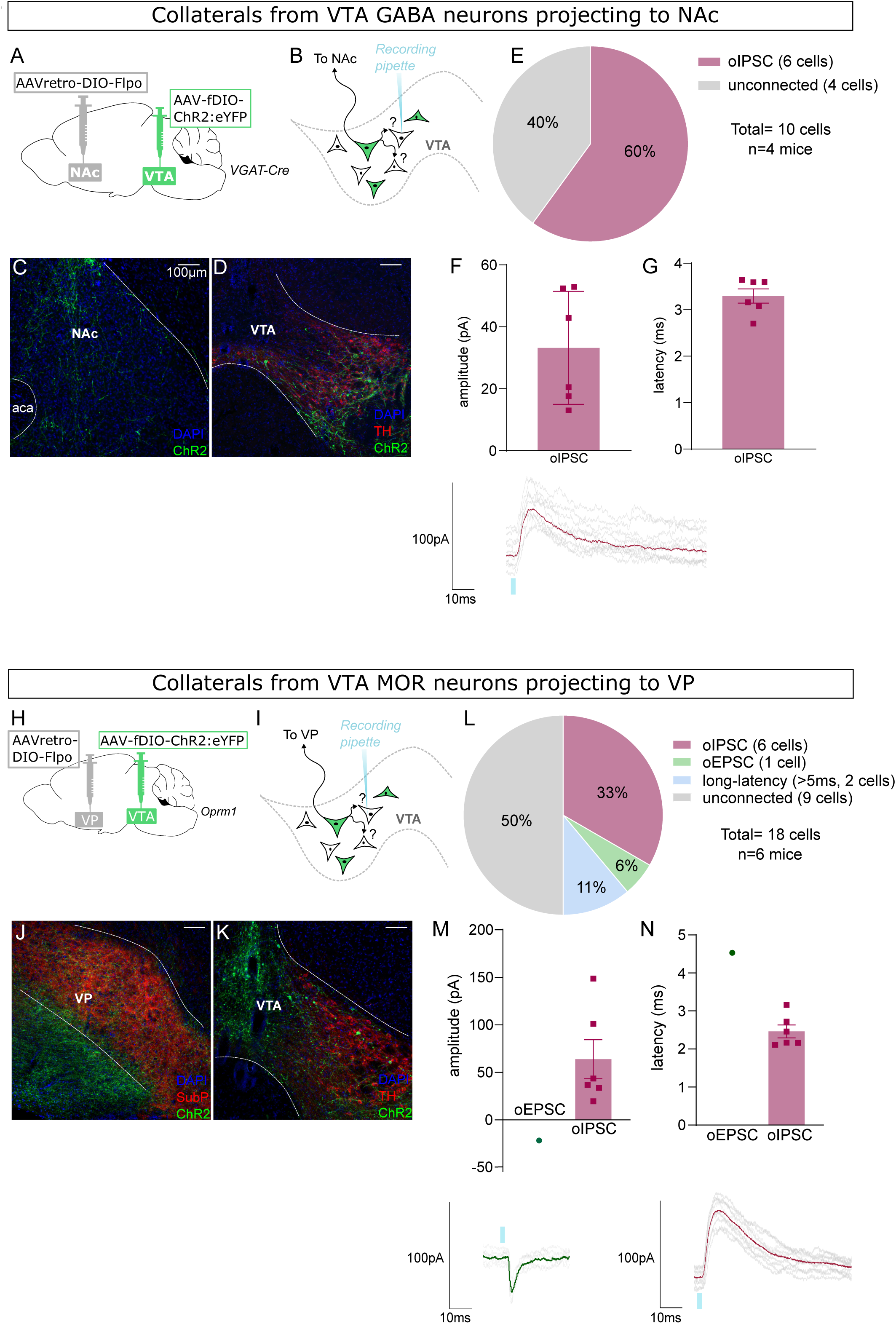
NAc-projecting VTA GABA neurons, and VP-projecting VTA MOR neurons, make intra-VTA synapses. (**A**) Dual AAV approach to express ChR2:eYFP in NAc-projecting VTA neurons in VGAT-Cre mice. (**B**) Patch-clamp recordings from ChR2:eYFP-negative neurons of VTA to test for collateralizing synapses made by NAc-projectors. (**C**) Coronal images showing ChR2:eYFP expression in NAc and (**D**) VTA; scale bars 100 µm. (**E**) ChR2:eYFP-negative VTA neuron responses to optogenetic stimulation of NAc-projectors. (**F**) Peak amplitude of connected cells that displayed an oIPSC, with example trace. (**G**) Latency to oIPSC onset. (**H**) Dual AAV approach to express ChR2:eYFP in VP-projecting VTA neurons in MOR-Cre mice. (**I**) Patch-clamp recordings from ChR2:eYFP-negative neurons of VTA to test for collateralizing synapses made by VP-projectors. (**J**) Coronal images showing ChR2:eYFP expression in VP and (**K**) VTA; scale bars 100 µm. (**L**) ChR2:eYFP-negative VTA neuron responses to optogenetic stimulation of VP-projectors. (**M**) Peak amplitude of connected cells that displayed an oEPSC and/or oIPSC (excluding long-latency), with example traces. (**N**) Latency to oPSC onset (excluding long-latency).

Finally, we did a similar experiment using MOR-Cre mice to target VP-projecting VTA neurons (**Figure 7H-K**). Here we found that half of the recorded YFP-negative cells were connected, showing primarily short-latency oIPSCs along with fewer short- and long-latency oEPSCs (**Figure 7L-N**).

## DISCUSSION

The VTA plays consequential roles in the orchestration of motivated behaviors and is composed of heterogeneous populations of DA, GABA, and glutamate neurons that send dense projections to diverse forebrain regions (Fields et al., 2007; Morales & Margolis, 2017). Yet VTA DA, GABA, and glutamate neurons also release their neurotransmitters locally within VTA. DA is released from somatodendritic compartments, activating DA autoreceptors (Ford, 2014). Multiple lines of evidence indicate that GABA and glutamate neurons resident to VTA synapse locally on to VTA DA and non-DA neurons (Beier, 2022; Dobi et al., 2010; Omelchenko & Sesack, 2009; Soden et al., 2020; Tan et al., 2012). VTA GABA neurons that make local synapses within VTA have frequently been described as interneurons (Bonci & Williams, 1996; Johnson & North, 1992; Luscher & Malenka, 2011; O’Brien & White, 1987). But there is scant evidence that VTA GABA interneurons and projection neurons represent distinct cell types. In this study we used retroAAV to target VTA neurons that project to NAc, VP, PFC, or LHb for recombinase-dependent expression of synaptic tags to image putative release sites, or of opsin for optogenetic stimulation of projection neurons while recording synaptic events in neighboring VTA neurons. Both approaches point to the same conclusion, that at least a subset of GABA and glutamate projection neurons collateralize locally and make intra-VTA synapses.

In cortex, hippocampus and other areas dominated by glutamate projection neurons the term interneuron is often used to describe inhibitory GABA neurons that synapse on to neurons within the same structure as their soma reside. However, the limbic basal ganglia circuits in which VTA neurons are embedded include many GABAergic projection neurons, at least subsets of which are understood to make both distal and local synapses. For example, DA D2 receptor-expressing medium spiny neurons are GABA projection neurons that also make extensive local collaterals that laterally inhibit and regulate other striatal neurons (Dobbs et al., 2016; Tunstall et al., 2002). Thus, a meaningful definition of the term in the context of mesolimbic circuitry, and the definition of interneuron we use in this study, is a neuron that synapses locally but not distally. Indeed, this definition captures many well characterized populations of neurons throughout the cortex, hippocampus, striatum, or cerebellum (Lim et al., 2018; Maccaferri & Lacaille, 2003; Pelkey et al., 2017; Petilla Interneuron Nomenclature et al., 2008). For example, cortical or striatal interneurons that express PV, SST, or cholinergic markers can be readily distinguished at the molecular level, but also by physiological properties that distinguish them from projection neurons or other cell types (Huang & Paul, 2019; Markram et al., 2004; Tepper et al., 2018).

The VTA has been known to contain non-DA GABA neurons since at least the early 1980’s (Gysling & Wang, 1983; Oertel & Mugnaini, 1984; Waszczak & Walters, 1980; Yim & Mogenson, 1980). Moreover, VTA GABA neurons were demonstrated to make inhibitory synapses on to VTA DA neurons (Bayer & Pickel, 1991; Johnson & North, 1992). VTA DA neurons may be distinguished from non-DA neurons (at least in lateral VTA) based on firing rate and other physiological and pharmacological features (Bunney et al., 1973; German et al., 1980; Waszczak & Walters, 1980). These observations serve as the primary basis for the notion of a VTA interneuron. However, those observations could instead be explained by VTA GABA projection neurons that collateralize locally. Indeed, one notable study used in vivo electrophysiology to identify a population of non-DA projection neurons and showed that they were reliably activated by antidromic stimulation of the internal capsule (Steffensen et al., 1998). This suggests that the population of non-DA VTA neurons they were able to identify through in vivo recordings were projection neurons. Likewise, substantia nigra (SN) compacta DA neurons were inhibited by antidromically identified GABA projection neurons in SN reticulata (Tepper et al., 1995), suggesting a parallel between our findings in VTA and those in SN.

If GABA interneurons represent one or more bona fide VTA cell types, then it is reasonable to suppose that they would be distinguishable by a molecular marker (or a constellation of markers). Markers that confer a GABAergic identity label VTA GABA projection neurons, and thus could not distinguish putative interneurons from projection neurons. However, several other markers have been shown to co-localize with a subset of VTA GABA but not DA neurons and thus represent potential interneuron markers (Bouarab et al., 2019; Nagaeva et al., 2020; Olson & Nestler, 2007; Phillips et al., 2022). We selected four of these markers that had well-validated Cre lines: PV, SST, MOR, and NTS. Using two different tracing strategies we confirmed that these markers are expressed in a subset of VTA neurons that are primarily non-dopaminergic. We found that PV^+^ VTA neurons project densely to LHb (and weakly to other projection targets), while SST^+^, NTS^+^, and especially MOR^+^ VTA neurons project densely to VP (and other projection targets).

Thus, while it is possible that there exists VTA interneurons that express one or more of these markers, none of these markers can be used on its own to discriminate between VTA projection neurons and VTA interneurons.

While the retroAAV approach resulted in strong labeling of projection neurons, including GABA- and glutamate-releasing neurons, it is likely that the intrinsic tropism of retroAAV influenced the population of projection cells that we labeled. Indeed, prior work showed that midbrain DA neurons are not efficiently targeted by the retroAAV vector we used (Tervo et al., 2016). Thus, populations of neurons that release DA and co-release GABA or glutamate may not contribute to the signals we measured.

In addition to GABA projection neurons, our experiments revealed that glutamate projection neurons in VTA also make local synapses. Therefore, the local excitatory synaptic events observed in prior studies (Dobi et al., 2010; McGovern et al., 2023; Yoo et al., 2016) may be driven by collaterals made by VTA glutamate projection neurons rather than a population of glutamate interneurons. Interestingly, we found that optogenetic activation of unspecified VTA projection neurons induced intra-VTA oEPSCs more rarely than intra-VTA oIPSCs. However, we also observed long-latency oPSCs, that were likely the result of activating VTA glutamate projection neurons that make intra-VTA excitatory collaterals and drive feed-forward recruitment of other VTA cells that also make local synapses.

The VTA integrates a large number of inhibitory inputs from a multitude of brain regions. However, recent studies indicate that neurons local to VTA preferentially inhibit DA neurons compared to GABAergic afferents from distal sources (Beier, 2022; Soden et al., 2020). Yet these studies cannot determine whether the local neurons synapsing on to VTA DA neurons are interneurons versus collaterals made by projection neurons. Moreover, VTA has been shown to receive an important GABAergic input from the rostral medial tegmental nucleus (RMTg) (Jhou, 2005; Perrotti et al., 2005). While VTA and RMTg are considered separate structures, the boundary between caudal VTA and rostral RMTg is ambiguous, and this area is dominated by GABA neurons (Smith et al., 2019). Interestingly, RMTg inhibitory synapses on to VTA DA neurons are strongly inhibited by MOR activation, and thus some of the functions classically attributed to VTA interneurons may be mediated by these short-range projection neurons (Jhou, 2021; Jhou et al., 2009; Kaufling & Aston-Jones, 2015; Matsui et al., 2014; St Laurent et al., 2020).

Within VTA, we found that MOR and several other potential interneuron markers were instead expressed in projection neurons. Other interneuron marker candidates have been suggested but, to our knowledge, no other VTA marker has been shown to be expressed selectively within a VTA interneuron population (Bouarab et al., 2019; Paul et al., 2019). One promising candidate is neuronal nitric oxide synthase (nNOS) which labels a subset of VTA GABA neurons in the parabrachial pigmented area of VTA that may not project distally, but also labels DA and glutamate neurons in adjacent areas of VTA and substantia nigra (Paul et al., 2018). Future work, for example using intersectional labeling, may resolve whether nNOS selectively labels bona fide GABA interneurons in VTA. Another candidate marker of interest is prepronociceptin (PNOC). A recent report showed that PNOC labels a population of paranigral non-DA neurons that make dense intra-VTA synapses without projecting to NAc (Parker et al., 2019). However, VTA PNOC neurons express both GABA and glutamate markers, and it is not clear whether they project to other VTA projection targets, such as VP or LHb.

In sum, we provide multiple lines of evidence that VTA GABA (and glutamate) neurons that project to distal targets also collateralize locally and make intra-VTA synapses. We also demonstrate that several candidate markers, including MOR, are expressed in VTA projection neurons. Future efforts may reveal positive evidence for the existence of VTA interneurons, for example through the identification of a marker, or a combinatorial set of markers, that labels VTA neurons that make local but not distal connections. At present, however, there is little evidence to support the notion of a VTA interneuron. We suggest that some functions prior attributed to VTA interneurons, such as MOR-mediated disinhibition of DA neurons, may instead be mediated by VTA projection neurons that make synaptic collaterals on to DA neurons. In this way the actions of opioids on VTA neurons would not only disinhibit DA neurons, but simultaneously inhibit GABA (or glutamate) release from distal VTA projections to VP and elsewhere. Indeed, in light of our increasing understanding for the roles of VTA GABA and glutamate projections in processes underlying behavioral reinforcement, their direct effects on distal targets may contribute to opioid-induced behaviors or adaptations relevant to drug addiction distinct from their effects on VTA DA neurons.

## METHODS

### Animals

Mice were group-housed (up to 5 mice/cage), bred at the University of California, San Diego (UCSD), kept on a 12h light-dark cycle, and had access to food and water ad libitum. Initial breeders were acquired from The Jackson Laboratory (**Table 1**), except for the MOR-Cre (Bailly et al., 2020) obtained from the lab of Brigitte Kieffer (University of Strasbourg). All mice were bred with a C57Bl/6 background and used as a mix of heterozygotes and homozygotes). Male and female mice were used in all experiments. All experiments were performed in accordance with protocols approved by the UCSD Institutional Animal Care and Use Committee.

**Table 1.**
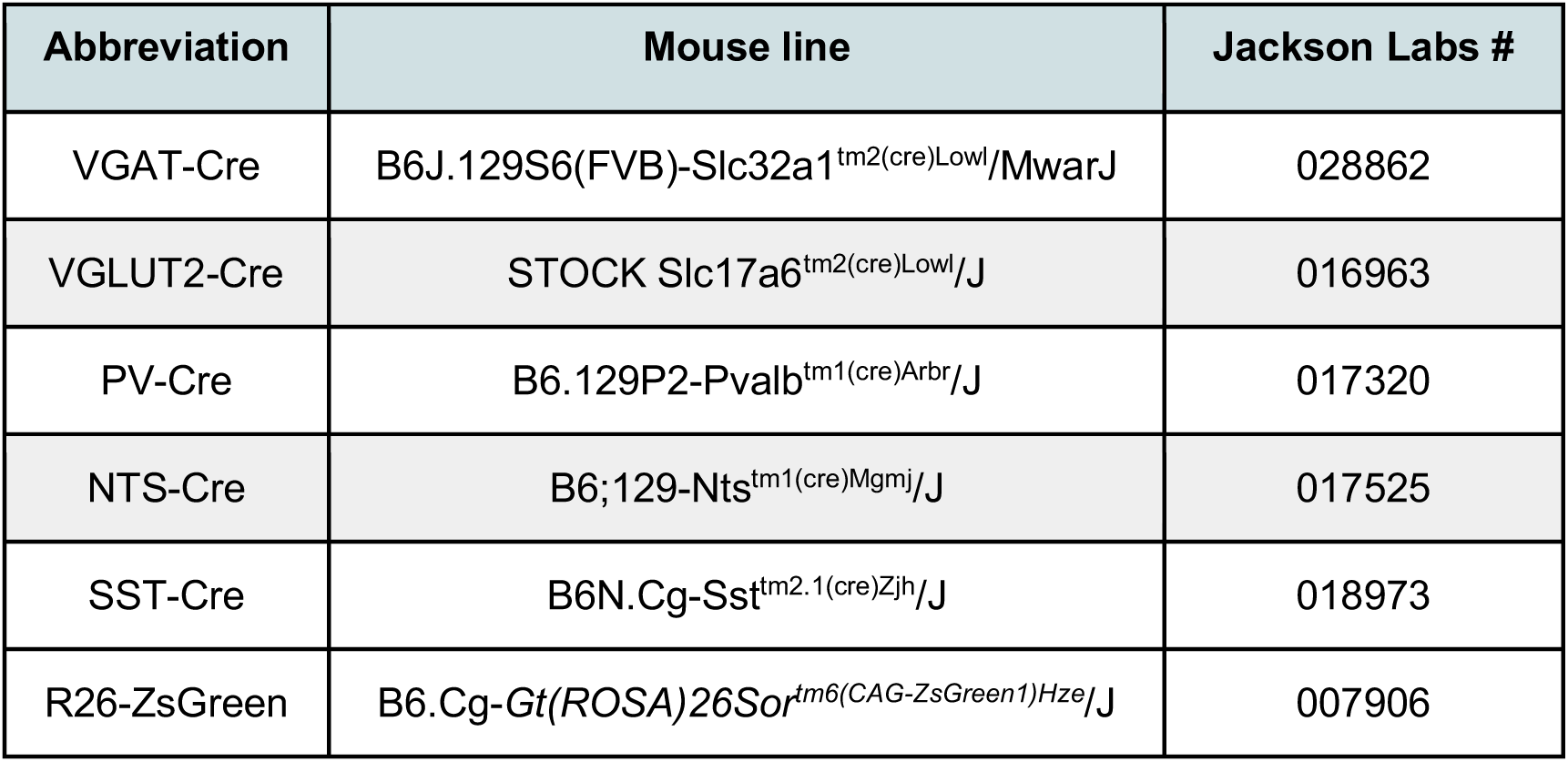
Mouse lines.

### Stereotaxic surgery

Mice >5 weeks (and up to 6months old) were deeply anesthetized with Isoflurane (502017, Primal Critical Care) and placed on a stereotaxic frame (Kopf 1900) for microinjection into discrete brain areas (**Table 2**). After ensuring the skull is flat small holes were drilled (1911-C Kopf) and AAVs (**Table 3**) infused with Nanoject (3-000-207, Drummond) using glass injectors (3-000-203-G/X, Drummond) pulled on a horizontal pipette puller (P-1000 Sutter Instrument). After infusion the injector was left for 3 to 5 min then withdrawn. Analgesia was provided via injections with 5mg/kg S.C. Carprofen (510510 Vet One). Electrophysiology was performed >3 weeks after surgery, histology >5 weeks.

**Table 2.** Stereotaxic coordinates.

| Injection site | ML | AP | DV |
| --- | --- | --- | --- |
| VTA | -0.35 | -3.35 | -4.3 |
| NAc | -0.8 | 1.34 | -4.5 |
| PFC | -0.4 | +1.9 | -1.7 |
| VP | -1.45 | +0.55 | -5.35 |
| VTA (MOR-Cre) | -0.6 | -.3.4 | -4.4 |

**Table 3.** AAV vectors.

| AAV | Titer | Packaged by | Volume | Addgene # |
| --- | --- | --- | --- | --- |
| AAVretro-EF1a-Cre | 3*10<sup>13</sup> | Salk GT3 | 150nL | 55636 |
| AAV5-EF1α-DIO-hChR2(H134R)-mCherry | 2*10<sup>13</sup> | Addgene | 150nL for ephys<br>100nL for histology | 20297 |
| AAVDJ-hSyn1-FLEXFRT mGFP-2A-Synaptophysin:mRuby | 2*10<sup>13</sup> | Addgene | 150nL | 71761 |
| AAVretro-hSyn1-DIO-Flpo | 2*10<sup>12</sup> | Salk GT3 | 150nL | NA |

### Histology

Mice were deeply anesthetized with pentobarbital (200 mg.kg-1, i.p., 200-071, Virbac) and transcardially perfused with 30mL of PBS (BP399, Fisher bioreagents) followed by 50mL of 4% PFA (18210, Electron Microscopy Sciences) in PBS. Brains were removed, post-fixed in 4% PFA overnight, and dehydrated in 30% sucrose (S0389, Sigma-Aldrich) in PBS for 48h then flash-frozen in isopentane. Brains were cut in 30µm coronal sections on a cryostat (CM3050S, Leica). Sections were selected to encompass the VTA and efferents to PFC, NAc, VP, and LHb. Sections were blocked in 5% normal donkey serum/0.4% Triton X-100 in PBS for 1h at room temperature and incubated with primary antibodies (**Table 4**) overnight at 4°C in the blocking buffer. Next day, slides were washed three times in 0.4% Triton X-100 in PBS for 5 min and incubated with secondary antibodies for two hours at room temperature shielded from the light. Finally, sections were washed three times in 0.4% Triton X-100 in PBS for 5 min and coverslipped with Fluoromount-G (Southern Biotech) containing 0.5µg/mL of DAPI (Roche). Images were taken using a Zeiss Axio Observer Epifluorescence microscope. For Supplemental Figure 1, the same procedure was used but the brains were cut sagittaly at a 15° angle.

**Table 4.**
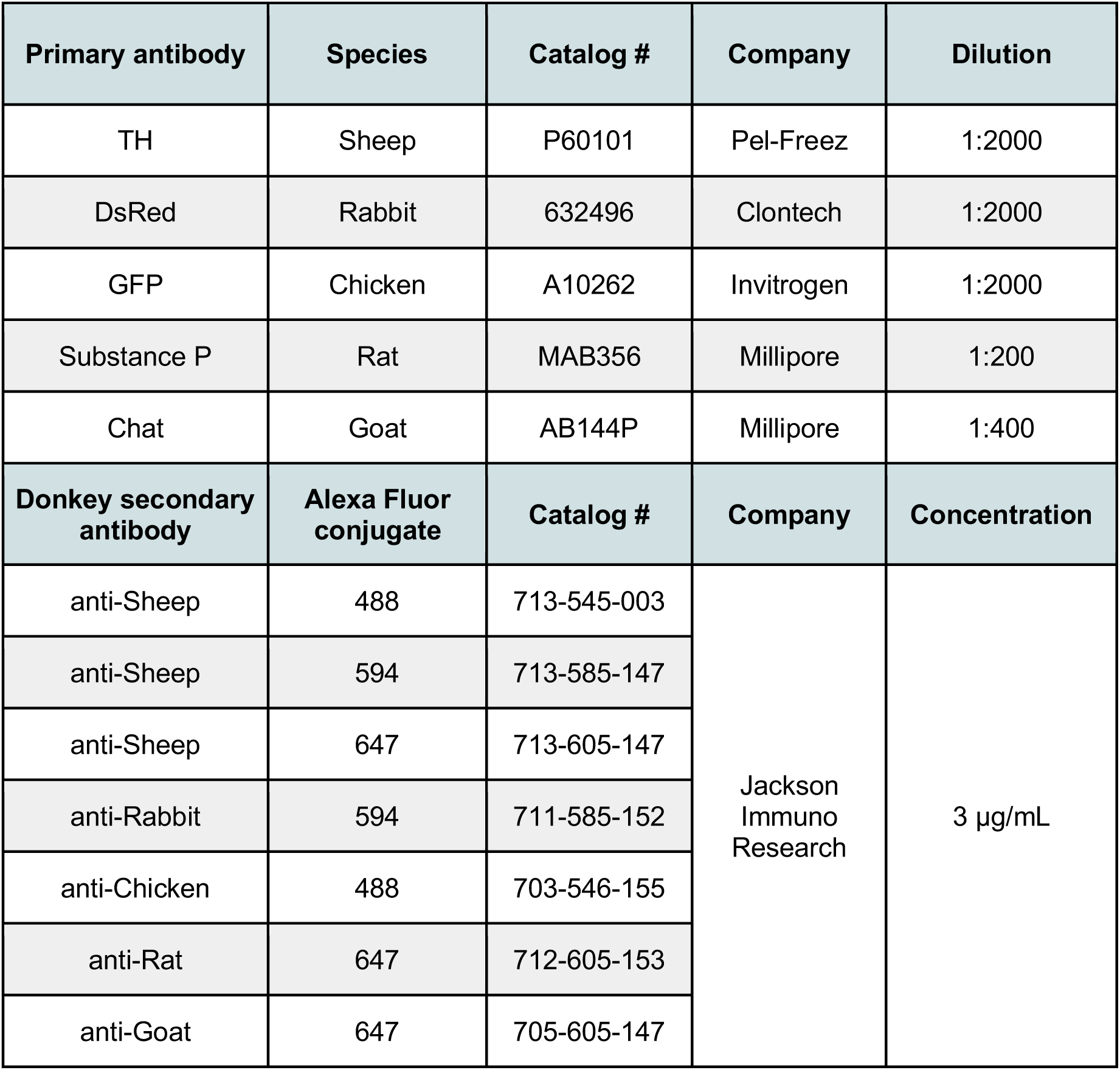
Antibodies.

| Primary antibody | Species | Catalog # | Company | Dilution |
| --- | --- | --- | --- | --- |
| TH | Sheep | P60101 | Pel-Freez | 1:2000 |
| DsRed | Rabbit | 632496 | Clontech | 1:2000 |
| GFP | Chicken | A10262 | Invitrogen | 1:2000 |
| Substance P | Rat | MAB356 | Millipore | 1:200 |
| Chat | Goat | AB144P | Millipore | 1:400 |
| Donkey secondary antibody | Alexa Fluor conjugate | Catalog # | Company | Concentration |
| anti-Sheep | 488 | 713-545-003 | Jackson Immuno Research | 3 µg/mL |
| anti-Sheep | 594 | 713-585-147 |  |  |
| anti-Sheep | 647 | 713-605-147 |  |  |
| anti-Rabbit | 594 | 711-585-152 |  |  |
| anti-Chicken | 488 | 703-546-155 |  |  |
| anti-Rat | 647 | 712-605-153 |  |  |
| anti-Goat | 647 | 705-605-147 |  |  |

### Colocalization with TH and counting

For each genetic marker, 3-4 mice and 4 sections through VTA per mouse were stained with antibodies against TH and DsRed. All sections were imaged at 10X with the same exposure parameters, using a Zeiss AxioObserver equipped with Apotome2 for structured illumination. The same display settings were applied to all images within condition. TH signal was used to define the boundaries of VTA and align to Bregma point. Cells expressing mCherry were identified first, then scored for presence or absence of TH expression. The counts were done independently by two experimenters and a high correlation was observed between the experimenters (R^2^=0.77, p<0.001, 60 total sections). Each cell that was only identified by one observer was reassessed for inclusion in final dataset.

### Single injection tracing

For evaluation of projection targets following a single AAV injection into VTA, we excluded subjects that had <30% of labeled cell bodies outside the VTA (**Table 5**). We also excluded subjects that had mCherry-labeled cell bodies in supramammillary nucleus. But we did not exclude mice with spread to red nucleus or IPN because these regions are not known to project to NAc, PFC, VP, or LHb (Liang et al., 2011; McLaughlin et al., 2017).

**Table 5.**
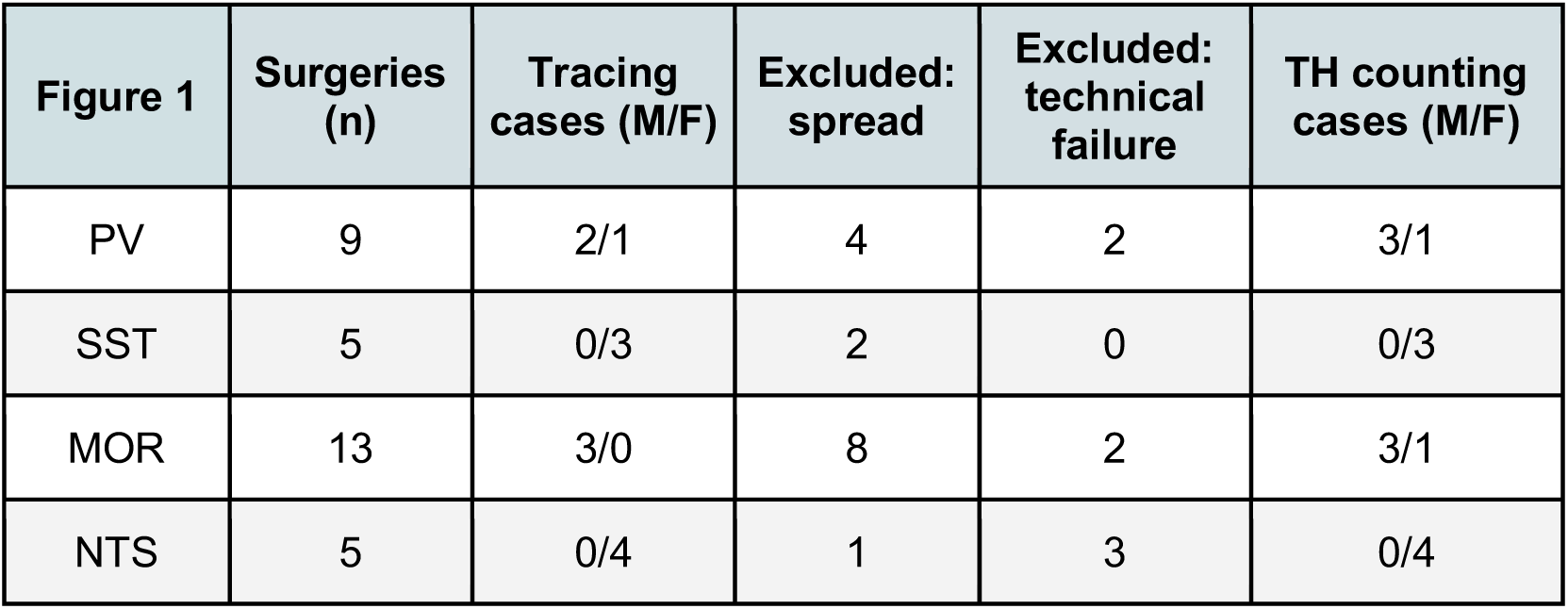
Cases included/excluded for Figure 1.

### Electrophysiology

Mice were deeply anesthetized using pentobarbital (200 mg/kg, i.p., Virbac) and transcardially perfused with 30 ml cold N-methyl D-glucamine (NMDG)-artificial Crebro-Spinal Fluid (aCSF, containing in mM: 92 NMDG, 2.5 KCl, 1.25 NaH2PO4, 30 NaHCO3, 20 HEPES, 25 D-glucose, 5 sodium ascorbate, 2 thiourea, 3 sodium pyruvate, 10 MgSO4, 0.5 CaCl2, pH 7.3) saturated with carbogen (95% O2–5% CO2). Sections (coronal, 200µm) were cut through VTA while immersed in cold NMDG-aCSF using a vibratome (VT1200S, Leica). Slices were incubated at 33°C for 25-30 min in a holding chamber containing NMDG-aCSF saturated with carbogen. During the incubation NaCl concentration was slowly increased in 5-min increments by spiking the holding-aCSF with a 2M NaCl solution diluted with the NMDG-aCSF (Ting et al., 2018). Slices were incubated at 25°C for 30-45 min in a holding chamber containing holding-aCSF (containing in mM: 115 NaCl, 2.5 KCl, 1.23 NaH2PO4, 26 NaHCO3, 10 D-glucose, 5 sodium ascorbate, 2 Thiourea, 3 sodium pyruvate, 2 MgSO4, 2 CaCl2, pH7.3) saturated with carbogen. While recording, slices were superfused with 31°C recording-aCSF (containing in mM: 125 NaCl, 2.5 KCl, 1.20 NaH2PO4, 26 NaHCO3, 12.5 D-glucose, 2 MgSO4, 2 CaCl2) using an in-line heater (TC-324B, Warner) at 1.5 ml/min. Whole-cell patch-clamp recordings from mCherry-negative VTA neurons were performed under visual guidance with infrared illumination and differential interference contrast using a Zeiss Axiocam MRm, Examiner.A1 equipped with a 40X objective. 6-7MΩ patch pipettes were pulled from borosilicate glass (Sutter Instruments) and filled with internal solution (containing in mM: 133.4 cesium-methanesulfonate, 22.7 HEPES, 0.45 EGTA, 3.2 NaCl, 5.7 tetraethylammonium-chloride, 0.48 NA-GTP, 4.5 Na2-ATP, pH to 7.3 with Cesium-OH). Postsynaptic currents were recorded in whole-cell voltage clamp (Multiclamp 700B amplifier, Axon Instruments), filtered at 2 kHz, digitized at 20 kHz (Axon Digidata 1550, Axon Instruments), and collected using pClamp 10 software (Molecular Device). Neurons were first held at -65mV to record excitatory currents and then at 0mV to record inhibitory currents. Optogenetic-evoked postsynaptic currents (oPSCs) were induced by flashing blue light (two 10Hz 2ms pulses, every 15s) through the light path of the microscope using a light-emitting diode (UHP-LED460, Prizmatix, 50mW) under computer control. We discarded likely ChR2+ cells, displaying photocurrent (**Supplemental Figure 5**), identified as starting within 1 ms of the light pulse, as well as cells where the series resistance varied by more than 20%. After breaking-in we waited 2-3 minutes before beginning optogenetic stimulation. For each cell we first recorded a baseline period (4-6 min) and for some cells baseline was followed by 4-6 min bath application of drug: 1µM TTX, 50-100µM 4AP, 10µM DNQX, 100µM PTX (**Table 6**). For each condition we averaged the last 10 sweeps; amplitude represented the peak current, and latency calculated as the duration from light onset to current onset.

**Table 6.** Drugs and physiology reagents.

| Reagent | Catalog # | Company |
| --- | --- | --- |
| 4AP | 0940 | Tocris |
| CaCl<sub>2</sub>·2H<sub>2</sub>O | BP510 | Fisher bioreagents |
| Ces met | 2550-61-0 | Sigma-Aldrich |
| D-Glucose | G8270 | Sigma |
| DNQX | D0540 | Sigma |
| EGTA | E3889 | Sigma |
| HEPES | H3375 | Sigma |
| KCl | BP366 | Fisher bioreagents |
| MgSO<sub>4</sub>·7H<sub>2</sub>O | M80 | Fisher bioreagents |
| Na-GTP | G8877 | Sigma |
| Na<sub>2</sub>-ATP | A2383 | Sigma |
| NaCl | BP358 | Fisher bioreagents |
| NaH<sub>2</sub>PO<sub>4</sub> | BP329 | Fisher bioreagents |
| NaHCO<sub>3</sub> | BP328 | Fisher bioreagents |
| NMDG | M2004 | Sigma-Aldrich |
| PTX | P1675 | Sigma |
| Sodium ascorbate | A7631 | Sigma |
| Sodium pyruvate | P2256 | Sigma-Aldrich |
| TEA chloride | 86616 | Fluka |
| Thiourea | T8656 | Sigma-Aldrich |
| TTX | 1069 | Tocris |

### Statistics

Data values are presented as means ± SEM. Effects of drug application were subjected to Friedman’s test (nonparametric ANOVA) followed by a Dunn’s posthoc test. Statistical significance was set at p<0.05.

## Author Contributions

Lucie Oriol: <u>Contribution,</u> methodology, investigation (stereotaxic surgeries, histology, electrophysiology, microscopy, cell counting), analysis and validation, visualization, project coordination and writing. <u>Competing interests,</u> no competing interests declared

Melody Chao: <u>Contribution,</u> investigation (histology) and visualization. <u>Competing interests,</u> no competing interests declared.

Grace J. Kollman: <u>Contribution,</u> investigation (cell counting) and resources (breeding). <u>Competing interests</u>, no competing interests declared.

Dina S. Dowlat: <u>Contribution,</u> investigation (LHb stereotactic injections) and resources (reagents provision). <u>Competing interests</u>, no competing interests declared.

Sarthak M. Singhal: <u>Contribution,</u> investigation (Electrophysiology in VP). <u>Competing interests</u>, no competing interests declared.

Thomas Steinkellner: <u>Contribution</u>, investigation (cloning). <u>Competing interests,</u> no competing interests declared.

Thomas S. Hnasko: <u>Contribution,</u> conceptualization, methodology, supervision, funding acquisition, validation, resources, and writing. <u>Competing interests,</u> no competing interests declared.

## Acknowledgements

This work was supported by funds from the National Institutes of Health (R01DA036612) and Veterans Affairs (I01BX005782). We thank Karl Deisseroth and Liqun Luo for AAV vectors provided through Addgene (see methods), and Sarah Uran for technical assistance.

---

**Supplemental Figure 1:**
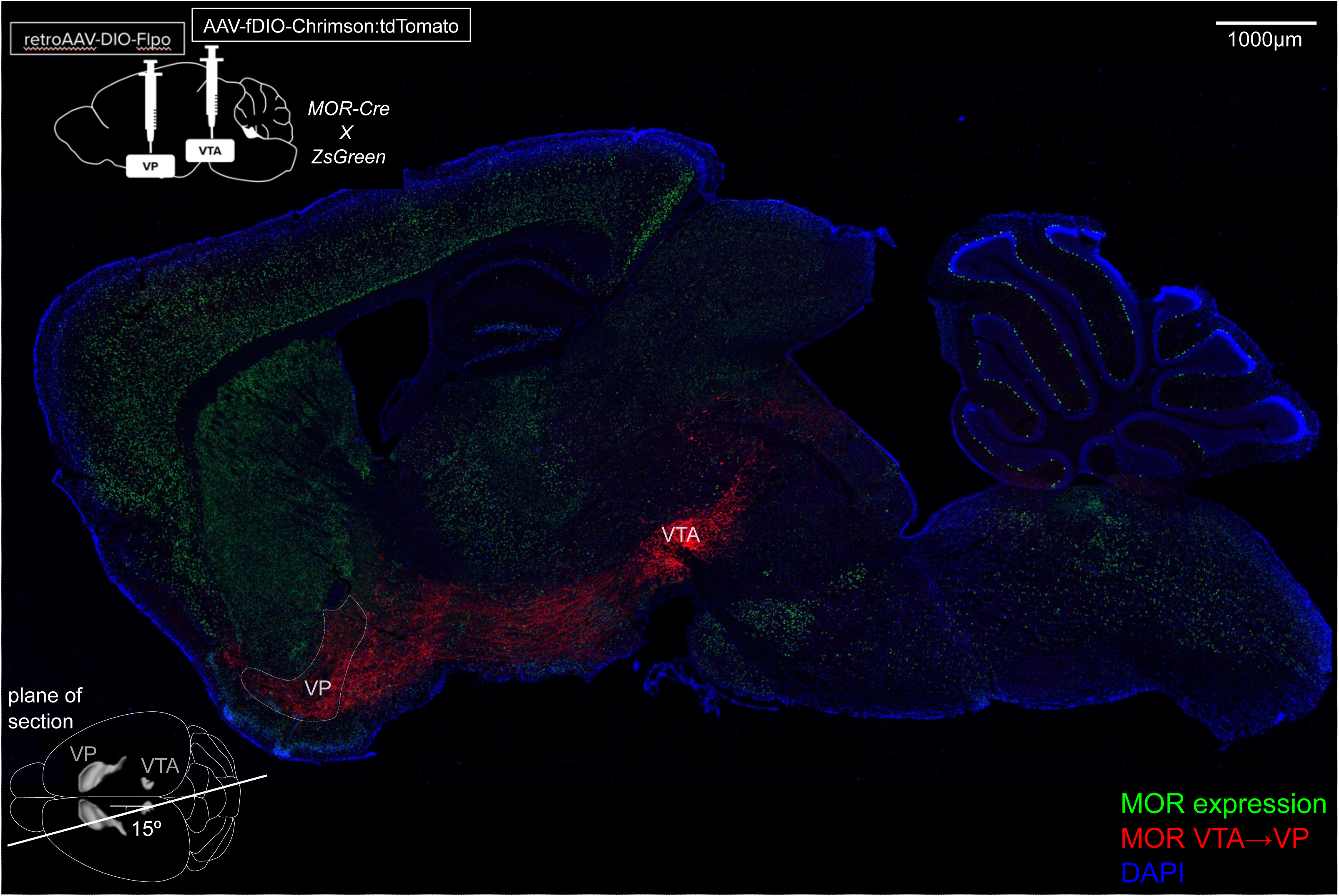
Projection of MOR-Cre-expressing VTA neurons to VP (related to Figure 3). Sagittal image, genotypes, and schematics of dual AAV approach and approximate location and sectioning angle of the cut. ZsGreen (green) labels all cells that have expressed MOR-Cre, Chrimson:tdTomato (red) labels cells/fibers from MOR-Cre VTA neurons projecting to VP, DAPI (blue) labels nuclei.

**Supplemental Figure 2:**
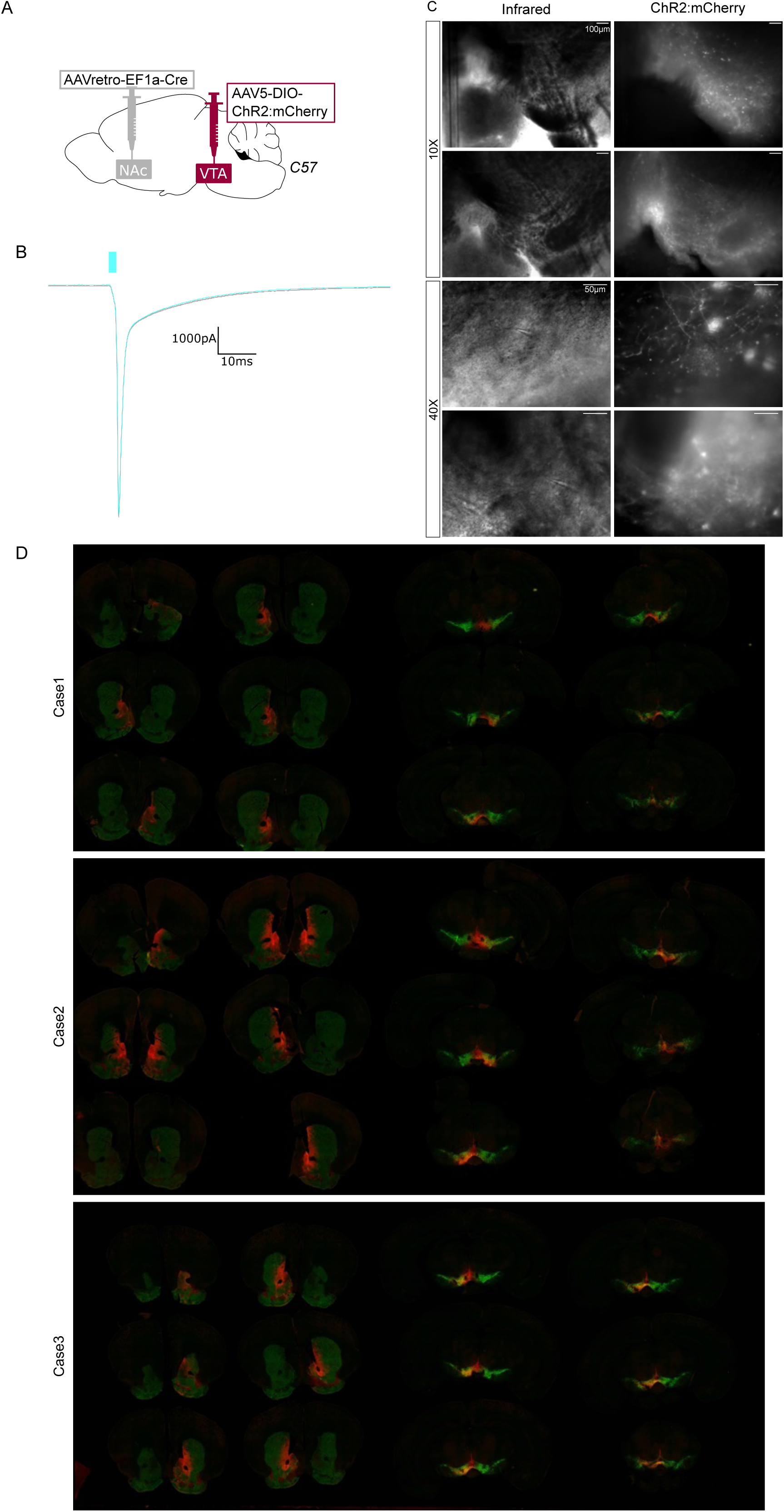
Photocurrent and histological validation of approach used in Figure 5 (related to Figure 5). (**A**) Dual AAV approach to express ChR2:mCherry in NAc-projecting VTA neurons in wild-type mice. (**B**) Example opsin-mediated photocurrent from ChR2:mCherry-positive neuron of VTA. (**C**) Example images under DIC IR light and mCherry expression around patch-clamp pipettes. (**D**) Additional cases of histology.

**Supplemental Figure 3:**
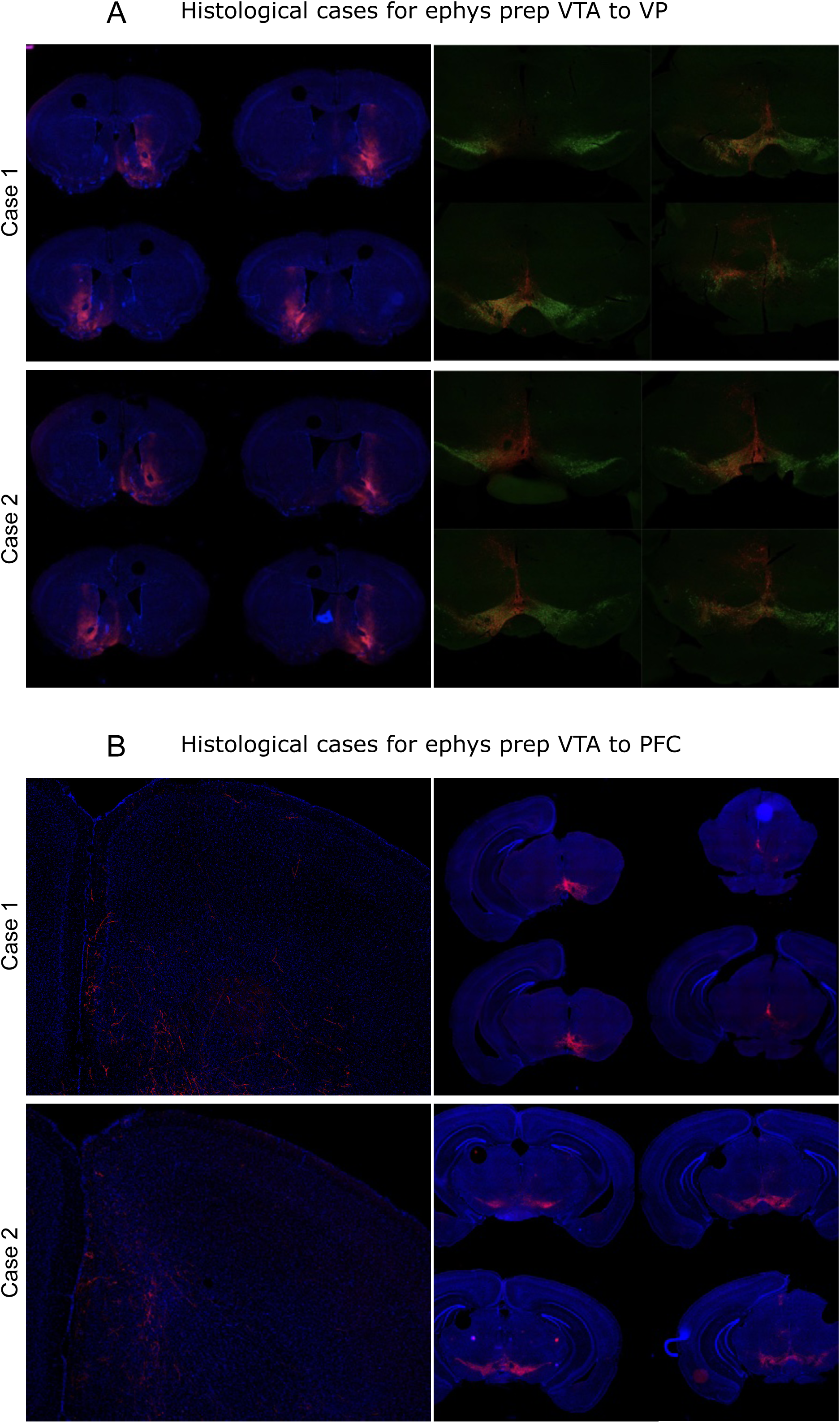
Histological validation of approach used in Figure 6 (related to Figure 6). (**A**) Additional cases of histology with expression of ChR2:mCherry in VP-projecting VTA neurons in wild-type mice. (**B**) Additional cases of histology with expression of ChR2:mCherry in PFC-projecting VTA neurons in wild-type mice.

## REFERENCES CITED

Azcorra, M., Gaertner, Z., Davidson, C., He, Q., Kim, H., Nagappan, S., Hayes, C. K., Ramakrishnan, C., Fenno, L., Kim, Y. S., Deisseroth, K., Longnecker, R., Awatramani, R., & Dombeck, D. A. (2023). Unique functional responses differentially map onto genetic subtypes of dopamine neurons. Nat Neurosci, 26(10), 1762–1774. 10.1038/s41593-023-01401-9

Badrinarayan, A., Wescott, S. A., Vander Weele, C. M., Saunders, B. T., Couturier, B. E., Maren, S., & Aragona, B. J. (2012). Aversive stimuli differentially modulate real-time dopamine transmission dynamics within the nucleus accumbens core and shell. J Neurosci, 32(45), 15779–15790. 10.1523/JNEUROSCI.3557-12.2012

Bailly, J., Del Rossi, N., Runtz, L., Li, J. J., Park, D., Scherrer, G., Tanti, A., Birling, M. C., Darcq, E., & Kieffer, B. L. (2020). Targeting Morphine-Responsive Neurons: Generation of a Knock-In Mouse Line Expressing Cre Recombinase from the Mu-Opioid Receptor Gene Locus. eNeuro, 7(3). 10.1523/ENEURO.0433-19.2020

Bayer, V. E., & Pickel, V. M. (1991). GABA-labeled terminals form proportionally more synapses with dopaminergic neurons containing low densities of tyrosine hydroxylase-immunoreactivity in rat ventral tegmental area. Brain Res, 559(1), 44–55. 10.1016/0006-8993(91)90285-4

Beier, K. (2022). Modified viral-genetic mapping reveals local and global connectivity relationships of ventral tegmental area dopamine cells. Elife, 11. 10.7554/eLife.76886

Bonci, A., & Williams, J. T. (1996). A common mechanism mediates long-term changes in synaptic transmission after chronic cocaine and morphine. Neuron, 16(3), 631–639. 10.1016/s0896-6273(00)80082-3

Bouarab, C., Thompson, B., & Polter, A. M. (2019). VTA GABA Neurons at the Interface of Stress and Reward. Front Neural Circuits, 13, 78. 10.3389/fncir.2019.00078

Bunney, B. S., Walters, J. R., Roth, R. H., & Aghajanian, G. K. (1973). Dopaminergic neurons: effect of antipsychotic drugs and amphetamine on single cell activity. J Pharmacol Exp Ther, 185(3), 560–571. https://www.ncbi.nlm.nih.gov/pubmed/4576427

Caceda, R., Kinkead, B., & Nemeroff, C. B. (2006). Neurotensin: role in psychiatric and neurological diseases. Peptides, 27(10), 2385–2404. 10.1016/j.peptides.2006.04.024

Conrad, W. S., Oriol, L., Kollman, G. J., Faget, L., & Hnasko, T. S. (2024). Proportion and distribution of neurotransmitter-defined cell types in the ventral tegmental area and substantia nigra pars compacta. Addiction Neuroscience, 13. 10.1016/j.addicn.2024.100183

Corre, J., van Zessen, R., Loureiro, M., Patriarchi, T., Tian, L., Pascoli, V., & Luscher, C. (2018). Dopamine neurons projecting to medial shell of the nucleus accumbens drive heroin reinforcement. Elife, 7. 10.7554/eLife.39945

de Jong, J. W., Afjei, S. A., Pollak Dorocic, I., Peck, J. R., Liu, C., Kim, C. K., Tian, L., Deisseroth, K., & Lammel, S. (2019). A Neural Circuit Mechanism for Encoding Aversive Stimuli in the Mesolimbic Dopamine System. Neuron, 101(1), 133–151 e137. 10.1016/j.neuron.2018.11.005

Dobbs, L. K., Kaplan, A. R., Lemos, J. C., Matsui, A., Rubinstein, M., & Alvarez, V. A. (2016). Dopamine Regulation of Lateral Inhibition between Striatal Neurons Gates the Stimulant Actions of Cocaine. Neuron, 90(5), 1100–1113. 10.1016/j.neuron.2016.04.031

Dobi, A., Margolis, E. B., Wang, H. L., Harvey, B. K., & Morales, M. (2010). Glutamatergic and nonglutamatergic neurons of the ventral tegmental area establish local synaptic contacts with dopaminergic and nondopaminergic neurons. J Neurosci, 30(1), 218–229. 10.1523/JNEUROSCI.3884-09.2010

Eshel, N., Bukwich, M., Rao, V., Hemmelder, V., Tian, J., & Uchida, N. (2015). Arithmetic and local circuitry underlying dopamine prediction errors. Nature, 525(7568), 243–246. 10.1038/nature14855

Faget, L., Oriol, L., Lee, W. C., Zell, V., Sargent, C., Flores, A., Hollon, N. G., Ramanathan, D., & Hnasko, T. S. (2024). Ventral pallidum GABA and glutamate neurons drive approach and avoidance through distinct modulation of VTA cell types. Nat Commun, 15(1), 4233. 10.1038/s41467-024-48340-y

Fields, H. L., Hjelmstad, G. O., Margolis, E. B., & Nicola, S. M. (2007). Ventral tegmental area neurons in learned appetitive behavior and positive reinforcement. Annu Rev Neurosci, 30, 289–316. 10.1146/annurev.neuro.30.051606.094341

Ford, C. P. (2014). The role of D2-autoreceptors in regulating dopamine neuron activity and transmission. Neuroscience, 282, 13–22. 10.1016/j.neuroscience.2014.01.025

German, D. C., Dalsass, M., & Kiser, R. S. (1980). Electrophysiological examination of the ventral tegmental (A10) area in the rat. Brain Res, 181(1), 191–197. 10.1016/0006-8993(80)91269-x

Gomez, J. A., Perkins, J. M., Beaudoin, G. M., Cook, N. B., Quraishi, S. A., Szoeke, E. A., Thangamani, K., Tschumi, C. W., Wanat, M. J., Maroof, A. M., Beckstead, M. J., Rosenberg, P. A., & Paladini, C. A. (2019). Ventral tegmental area astrocytes orchestrate avoidance and approach behavior. Nat Commun, 10(1), 1455. 10.1038/s41467-019-09131-y

Gorelova, N., Mulholland, P. J., Chandler, L. J., & Seamans, J. K. (2012). The glutamatergic component of the mesocortical pathway emanating from different subregions of the ventral midbrain. Cereb Cortex, 22(2), 327–336. 10.1093/cercor/bhr107

Gysling, & Wang. (1983). Morphine-induced activation of A10 dopamine neurons in the rat. 10.1016/0006-8993(83)90913-7

Hnasko, T. S., Hjelmstad, G. O., Fields, H. L., & Edwards, R. H. (2012). Ventral tegmental area glutamate neurons: electrophysiological properties and projections. J Neurosci, 32(43), 15076–15085. 10.1523/JNEUROSCI.3128-12.2012

Huang, Z. J., & Paul, A. (2019). The diversity of GABAergic neurons and neural communication elements. Nat Rev Neurosci, 20(9), 563–572. 10.1038/s41583-019-0195-4

Jhou, T. (2005). Neural mechanisms of freezing and passive aversive behaviors. J Comp Neurol, 493(1), 111–114. 10.1002/cne.20734

Jhou, T. C. (2021). The rostromedial tegmental (RMTg) “brake” on dopamine and behavior: A decade of progress but also much unfinished work. Neuropharmacology, 198, 108763. 10.1016/j.neuropharm.2021.108763

Jhou, T. C., Geisler, S., Marinelli, M., Degarmo, B. A., & Zahm, D. S. (2009). The mesopontine rostromedial tegmental nucleus: A structure targeted by the lateral habenula that projects to the ventral tegmental area of Tsai and substantia nigra compacta. J Comp Neurol, 513(6), 566–596. 10.1002/cne.21891

Johnson, S. W., & North, R. A. (1992). Opioids excite dopamine neurons by hyperpolarization of local interneurons. J Neurosci, 12(2), 483–488. 10.1523/JNEUROSCI.12-02-00483.1992

Kalivas, P. W. (1983). Behavioral and neurochemical effects of neurotesin microinjection into the VTA of the rat. Neuroscience, 8(3), 495–505.

Kaufling, J., & Aston-Jones, G. (2015). Persistent Adaptations in Afferents to Ventral Tegmental Dopamine Neurons after Opiate Withdrawal. J Neurosci, 35(28), 10290–10303. 10.1523/JNEUROSCI.0715-15.2015

Kaufling, J., Veinante, P., Pawlowski, S. A., Freund-Mercier, M. J., & Barrot, M. (2010). gamma-Aminobutyric acid cells with cocaine-induced DeltaFosB in the ventral tegmental area innervate mesolimbic neurons. Biol Psychiatry, 67(1), 88–92. 10.1016/j.biopsych.2009.08.001

Kawaguchi, Y. (1993). Physiological, morphological, and histochemical characterization of three classes of interneurons in rat neostriatum. J Neurosci, 13(11), 4908–4923. 10.1523/JNEUROSCI.13-11-04908.1993

Kawaguchi, Y., & Kondo, S. (2002). Parvalbumin, somatostatin and cholecystokinin as chemical markers for specific GABAergic interneuron types in the rat frontal cortex. J Neurocytol, 31(3-5), 277–287. 10.1023/a:1024126110356

Kawano, M., Kawasaki, A., Sakata-Haga, H., Fukui, Y., Kawano, H., Nogami, H., & Hisano, S. (2006). Particular subpopulations of midbrain and hypothalamic dopamine neurons express vesicular glutamate transporter 2 in the rat brain. J Comp Neurol, 498(5), 581–592. 10.1002/cne.21054

Keiflin, R., & Janak, P. H. (2015). Dopamine Prediction Errors in Reward Learning and Addiction: From Theory to Neural Circuitry. Neuron, 88(2), 247–263. 10.1016/j.neuron.2015.08.037

Lammel, S., Steinberg, E. E., Foldy, C., Wall, N. R., Beier, K., Luo, L., & Malenka, R. C. (2015). Diversity of transgenic mouse models for selective targeting of midbrain dopamine neurons. Neuron, 85(2), 429–438. 10.1016/j.neuron.2014.12.036

Liang, H., Paxinos, G., & Watson, C. (2011). Projections from the brain to the spinal cord in the mouse. Brain Struct Funct, 215(3-4), 159–186. 10.1007/s00429-010-0281-x

Lim, L., Mi, D., Llorca, A., & Marin, O. (2018). Development and Functional Diversification of Cortical Interneurons. Neuron, 100(2), 294–313. 10.1016/j.neuron.2018.10.009

Luscher, C. (2016). The Emergence of a Circuit Model for Addiction. Annu Rev Neurosci, 39, 257–276. 10.1146/annurev-neuro-070815-013920

Luscher, C., & Malenka, R. C. (2011). Drug-evoked synaptic plasticity in addiction: from molecular changes to circuit remodeling. Neuron, 69(4), 650–663. 10.1016/j.neuron.2011.01.017

Ma, S., Zhong, H., Liu, X., & Wang, L. (2023). Spatial Distribution of Neurons Expressing Single, Double, and Triple Molecular Characteristics of Glutamatergic, Dopaminergic, or GABAergic Neurons in the Mouse Ventral Tegmental Area. J Mol Neurosci, 73(6), 345–362. 10.1007/s12031-023-02121-2

Maccaferri, G., & Lacaille, J. C. (2003). Interneuron Diversity series: Hippocampal interneuron classifications--making things as simple as possible, not simpler. Trends Neurosci, 26(10), 564–571. 10.1016/j.tins.2003.08.002

Markram, H., Toledo-Rodriguez, M., Wang, Y., Gupta, A., Silberberg, G., & Wu, C. (2004). Interneurons of the neocortical inhibitory system. Nat Rev Neurosci, 5(10), 793–807. 10.1038/nrn1519

Matsui, A., Jarvie, B. C., Robinson, B. G., Hentges, S. T., & Williams, J. T. (2014). Separate GABA afferents to dopamine neurons mediate acute action of opioids, development of tolerance, and expression of withdrawal. Neuron, 82(6), 1346–1356. 10.1016/j.neuron.2014.04.030

McGovern, D. J., Polter, A. M., Prevost, E. D., Ly, A., McNulty, C. J., Rubinstein, B., & Root, D. H. (2023). Ventral tegmental area glutamate neurons establish a mu-opioid receptor gated circuit to mesolimbic dopamine neurons and regulate opioid-seeking behavior. Neuropsychopharmacology, 48(13), 1889–1900. 10.1038/s41386-023-01637-w

McLaughlin, I., Dani, J. A., & De Biasi, M. (2017). The medial habenula and interpeduncular nucleus circuitry is critical in addiction, anxiety, and mood regulation. J Neurochem, 142 Suppl 2(Suppl 2), 130-143. 10.1111/jnc.14008

Miranda-Barrientos, J., Chambers, I., Mongia, S., Liu, B., Wang, H. L., Mateo-Semidey, G. E., Margolis, E. B., Zhang, S., & Morales, M. (2021). Ventral tegmental area GABA, glutamate, and glutamate-GABA neurons are heterogeneous in their electrophysiological and pharmacological properties. Eur J Neurosci. 10.1111/ejn.15156

Morales, M., & Margolis, E. B. (2017). Ventral tegmental area: cellular heterogeneity, connectivity and behaviour. Nat Rev Neurosci, 18(2), 73–85. 10.1038/nrn.2016.165

Nagaeva, E., Zubarev, I., Bengtsson Gonzales, C., Forss, M., Nikouei, K., de Miguel, E., Elsila, L., Linden, A. M., Hjerling-Leffler, J., Augustine, G. J., & Korpi, E. R. (2020). Heterogeneous somatostatin-expressing neuron population in mouse ventral tegmental area. Elife, 9. 10.7554/eLife.59328

Nestler, E. J. (2005). Is there a common molecular pathway for addiction? Nat Neurosci, 8(11), 1445–1449. 10.1038/nn1578

Nieh, E. H., Matthews, G. A., Allsop, S. A., Presbrey, K. N., Leppla, C. A., Wichmann, R., Neve, R., Wildes, C. P., & Tye, K. M. (2015). Decoding neural circuits that control compulsive sucrose seeking. Cell, 160(3), 528–541. 10.1016/j.cell.2015.01.003

O’Brien, D. P., & White, F. J. (1987). Inhibition of non-dopamine cells in the ventral tegmental area by benzodiazepines: relationship to A10 dopamine cell activity. Eur J Pharmacol, 142(3), 343–354. 10.1016/0014-2999(87)90072-0

Oades, R. D., & Halliday, G. M. (1987). Ventral tegmental (A10) system: neurobiology. 1. Anatomy and connectivity. Brain Res, 434(2), 117–165. 10.1016/0165-0173(87)90011-7

Oertel, W. H., & Mugnaini, E. (1984). Immunocytochemical studies of GABAergic neurons in rat basal ganglia and their relations to other neuronal systems. Neurosci Lett, 47(3), 233–238. 10.1016/0304-3940(84)90519-6

Olson, V. G., & Nestler, E. J. (2007). Topographical organization of GABAergic neurons within the ventral tegmental area of the rat. Synapse, 61(2), 87–95. 10.1002/syn.20345

Omelchenko, N., & Sesack, S. R. (2009). Ultrastructural analysis of local collaterals of rat ventral tegmental area neurons: GABA phenotype and synapses onto dopamine and GABA cells. Synapse, 63(10), 895–906. 10.1002/syn.20668

Ostroumov, A., & Dani, J. A. (2018). Inhibitory Plasticity of Mesocorticolimbic Circuits in Addiction and Mental Illness. Trends Neurosci, 41(12), 898–910. 10.1016/j.tins.2018.07.014

Parker, K. E., Pedersen, C. E., Gomez, A. M., Spangler, S. M., Walicki, M. C., Feng, S. Y., Stewart, S. L., Otis, J. M., Al-Hasani, R., McCall, J. G., Sakers, K., Bhatti, D. L., Copits, B. A., Gereau, R. W., Jhou, T., Kash, T. J., Dougherty, J. D., Stuber, G. D., & Bruchas, M. R. (2019). A Paranigral VTA Nociceptin Circuit that Constrains Motivation for Reward. Cell, 178(3), 653–671 e619. 10.1016/j.cell.2019.06.034

Paul, E. J., Kalk, E., Tossell, K., Irvine, E. E., Franks, N. P., Wisden, W., Withers, D. J., Leiper, J., & Ungless, M. A. (2018). nNOS-Expressing Neurons in the Ventral Tegmental Area and Substantia Nigra Pars Compacta. eNeuro, 5(5). 10.1523/ENEURO.0381-18.2018

Paul, E. J., Tossell, K., & Ungless, M. A. (2019). Transcriptional profiling aligned with in situ expression image analysis reveals mosaically expressed molecular markers for GABA neuron sub-groups in the ventral tegmental area. Eur J Neurosci, 50(11), 3732–3749. 10.1111/ejn.14534

Pelkey, K. A., Chittajallu, R., Craig, M. T., Tricoire, L., Wester, J. C., & McBain, C. J. (2017). Hippocampal GABAergic Inhibitory Interneurons. Physiol Rev, 97(4), 1619–1747. 10.1152/physrev.00007.2017

Perrotti, L. I., Bolanos, C. A., Choi, K. H., Russo, S. J., Edwards, S., Ulery, P. G., Wallace, D. L., Self, D. W., Nestler, E. J., & Barrot, M. (2005). DeltaFosB accumulates in a GABAergic cell population in the posterior tail of the ventral tegmental area after psychostimulant treatment. Eur J Neurosci, 21(10), 2817–2824. 10.1111/j.1460-9568.2005.04110.x

Petilla Interneuron Nomenclature, G., Ascoli, G. A., Alonso-Nanclares, L., Anderson, S. A., Barrionuevo, G., Benavides-Piccione, R., Burkhalter, A., Buzsaki, G., Cauli, B., Defelipe, J., Fairen, A., Feldmeyer, D., Fishell, G., Fregnac, Y., Freund, T. F., Gardner, D., Gardner, E. P., Goldberg, J. H., Helmstaedter, M., … Yuste, R. (2008). Petilla terminology: nomenclature of features of GABAergic interneurons of the cerebral cortex. Nat Rev Neurosci, 9(7), 557–568. 10.1038/nrn2402

Phillips, R. A., 3rd, Tuscher, J. J., Black, S. L., Andraka, E., Fitzgerald, N. D., Ianov, L., & Day, J. J. (2022). An atlas of transcriptionally defined cell populations in the rat ventral tegmental area. Cell Rep, 39(1), 110616. 10.1016/j.celrep.2022.110616

Poulin, J. F., Gaertner, Z., Moreno-Ramos, O. A., & Awatramani, R. (2020). Classification of Midbrain Dopamine Neurons Using Single-Cell Gene Expression Profiling Approaches. Trends Neurosci, 43(3), 155–169. 10.1016/j.tins.2020.01.004

Roeper, J. (2013). Dissecting the diversity of midbrain dopamine neurons. Trends Neurosci, 36(6), 336–342. 10.1016/j.tins.2013.03.003

Root, D. H., Barker, D. J., Estrin, D. J., Miranda-Barrientos, J. A., Liu, B., Zhang, S., Wang, H. L., Vautier, F., Ramakrishnan, C., Kim, Y. S., Fenno, L., Deisseroth, K., & Morales, M. (2020). Distinct Signaling by Ventral Tegmental Area Glutamate, GABA, and Combinatorial Glutamate-GABA Neurons in Motivated Behavior. Cell Rep, 32(9), 108094. 10.1016/j.celrep.2020.108094

Root, D. H., Estrin, D. J., & Morales, M. (2018). Aversion or Salience Signaling by Ventral Tegmental Area Glutamate Neurons. iScience, 2, 51–62. 10.1016/j.isci.2018.03.008

Root, D. H., Mejias-Aponte, C. A., Qi, J., & Morales, M. (2014). Role of glutamatergic projections from ventral tegmental area to lateral habenula in aversive conditioning. J Neurosci, 34(42), 13906–13910. 10.1523/JNEUROSCI.2029-14.2014

Shields, A. K., Suarez, M., Wakabayashi, K. T., & Bass, C. E. (2021). Activation of VTA GABA neurons disrupts reward seeking by altering temporal processing. Behav Brain Res, 410, 113292. 10.1016/j.bbr.2021.113292

Smith, R. J., Vento, P. J., Chao, Y. S., Good, C. H., & Jhou, T. C. (2019). Gene expression and neurochemical characterization of the rostromedial tegmental nucleus (RMTg) in rats and mice. Brain Struct Funct, 224(1), 219–238. 10.1007/s00429-018-1761-7

Soden, M. E., Chung, A. S., Cuevas, B., Resnick, J. M., Awatramani, R., & Zweifel, L. S. (2020). Anatomic resolution of neurotransmitter-specific projections to the VTA reveals diversity of GABAergic inputs. Nat Neurosci, 23(8), 968–980. 10.1038/s41593-020-0657-z

Soden, M. E., Yee, J. X., & Zweifel, L. S. (2023). Circuit coordination of opposing neuropeptide and neurotransmitter signals. Nature, 619(7969), 332–337. 10.1038/s41586-023-06246-7

St Laurent, R., Martinez Damonte, V., Tsuda, A. C., & Kauer, J. A. (2020). Periaqueductal Gray and Rostromedial Tegmental Inhibitory Afferents to VTA Have Distinct Synaptic Plasticity and Opiate Sensitivity. Neuron, 106(4), 624–636 e624. 10.1016/j.neuron.2020.02.029

Stamatakis, A. M., Jennings, J. H., Ung, R. L., Blair, G. A., Weinberg, R. J., Neve, R. L., Boyce, F., Mattis, J., Ramakrishnan, C., Deisseroth, K., & Stuber, G. D. (2013). A unique population of ventral tegmental area neurons inhibits the lateral habenula to promote reward. Neuron, 80(4), 1039–1053. 10.1016/j.neuron.2013.08.023

Steffensen, S. C., Svingos, A. L., Pickel, V. M., & Henriksen, S. J. (1998). Electrophysiological characterization of GABAergic neurons in the ventral tegmental area. J Neurosci, 18(19), 8003–8015. 10.1523/JNEUROSCI.18-19-08003.1998

Tan, K. R., Yvon, C., Turiault, M., Mirzabekov, J. J., Doehner, J., Labouebe, G., Deisseroth, K., Tye, K. M., & Luscher, C. (2012). GABA neurons of the VTA drive conditioned place aversion. Neuron, 73(6), 1173–1183. 10.1016/j.neuron.2012.02.015

Taylor, S. R., Badurek, S., Dileone, R. J., Nashmi, R., Minichiello, L., & Picciotto, M. R. (2014). GABAergic and glutamatergic efferents of the mouse ventral tegmental area. J Comp Neurol, 522(14), 3308–3334. 10.1002/cne.23603

Tepper, J. M., Koos, T., Ibanez-Sandoval, O., Tecuapetla, F., Faust, T. W., & Assous, M. (2018). Heterogeneity and Diversity of Striatal GABAergic Interneurons: Update 2018. Front Neuroanat, 12, 91. 10.3389/fnana.2018.00091

Tepper, J. M., Martin, L. P., & Anderson, D. R. (1995). GABAA receptor-mediated inhibition of rat substantia nigra dopaminergic neurons by pars reticulata projection neurons. J Neurosci, 15(4), 3092–3103. 10.1523/JNEUROSCI.15-04-03092.1995

Tepper, J. M., Tecuapetla, F., Koos, T., & Ibanez-Sandoval, O. (2010). Heterogeneity and diversity of striatal GABAergic interneurons. Front Neuroanat, 4, 150. 10.3389/fnana.2010.00150

Tervo, D. G., Hwang, B. Y., Viswanathan, S., Gaj, T., Lavzin, M., Ritola, K. D., Lindo, S., Michael, S., Kuleshova, E., Ojala, D., Huang, C. C., Gerfen, C. R., Schiller, J., Dudman, J. T., Hantman, A. W., Looger, L. L., Schaffer, D. V., & Karpova, A. Y. (2016). A Designer AAV Variant Permits Efficient Retrograde Access to Projection Neurons. Neuron, 92(2), 372–382. 10.1016/j.neuron.2016.09.021

Ting, A. K. R., & van der Kooy, D. (2012). The neurobiology of opiate motivation. Cold Spring Harb Perspect Med, 2(10). 10.1101/cshperspect.a012096

Tunstall, M. J., Oorschot, D. E., Kean, A., & Wickensv, J. R. (2002). Inhibitory Interactions Between Spiny Projection Neurons in the Rat Striatum. J Neurophysiol, 1263–1269. 10.1152/jn.2002.88.3.1263

van Zessen, R., Phillips, J. L., Budygin, E. A., & Stuber, G. D. (2012). Activation of VTA GABA neurons disrupts reward consumption. Neuron, 73(6), 1184–1194. 10.1016/j.neuron.2012.02.016

Wang, H. L., Qi, J., Zhang, S., Wang, H., & Morales, M. (2015). Rewarding Effects of Optical Stimulation of Ventral Tegmental Area Glutamatergic Neurons. J Neurosci, 35(48), 15948–15954. 10.1523/JNEUROSCI.3428-15.2015

Warlow, S. M., Singhal, S. M., Hollon, N. G., Faget, L., Dowlat, D. S., Zell, V., Hunker, A. C., Zweifel, L. S., & Hnasko, T. S. (2024). Mesoaccumbal glutamate neurons drive reward via glutamate release but aversion via dopamine co-release. Neuron, 112(3), 488–499 e485. 10.1016/j.neuron.2023.11.002

Waszczak, B. L., & Walters, J. R. (1980). Intravenous GABA agonist administration stimulates firing of A10 dopaminergic neurons. Eur J Pharmacol, 66(1), 141–144. 10.1016/0014-2999(80)90308-8

Yamaguchi, T., Wang, H. L., Li, X., Ng, T. H., & Morales, M. (2011). Mesocorticolimbic glutamatergic pathway. J Neurosci, 31(23), 8476–8490. 10.1523/JNEUROSCI.1598-11.2011

Yim, C. Y., & Mogenson, G. J. (1980). Effect of picrotoxin and nipecotic acid on inhibitory response of dopaminergic neurons in the ventral tegmental area to stimulation of the nucleus accumbens. Brain Res, 199(2), 466–473. 10.1016/0006-8993(80)90705-2

Yoo, J. H., Zell, V., Gutierrez-Reed, N., Wu, J., Ressler, R., Shenasa, M. A., Johnson, A. B., Fife, K. H., Faget, L., & Hnasko, T. S. (2016). Ventral tegmental area glutamate neurons co-release GABA and promote positive reinforcement. Nat Commun, 7, 13697. 10.1038/ncomms13697

Zell, V., Steinkellner, T., Hollon, N. G., Warlow, S. M., Souter, E., Faget, L., Hunker, A. C., Jin, X., Zweifel, L. S., & Hnasko, T. S. (2020). VTA Glutamate Neuron Activity Drives Positive Reinforcement Absent Dopamine Co-release. Neuron, 107(5), 864–873 e864. 10.1016/j.neuron.2020.06.011

Zhou, W. L., Kim, K., Ali, F., Pittenger, S. T., Calarco, C. A., Mineur, Y. S., Ramakrishnan, C., Deisseroth, K., Kwan, A. C., & Picciotto, M. R. (2022). Activity of a direct VTA to ventral pallidum GABA pathway encodes unconditioned reward value and sustains motivation for reward. Sci Adv, 8(42), eabm5217. 10.1126/sciadv.abm5217

